# The Rossmann2 × 2 Fold Attains its Native Structure Via a Defined Pathway of Sequential and Cooperative Folding Units

**DOI:** 10.64898/2026.05.21.726993

**Authors:** Robert P. Sosa, Jeonghoon Kim, Alexander B. Tong, Sudeesh Krishnamurthy, Christina Volz, Bendix Heβ, Daniel Pineda, Carlos Bustamante

## Abstract

Despite progress in predicting protein structures, how proteins arrive at their native state remains a subject of continuous debate. We present a single molecule force spectroscopy study of conserved Rossmann2 × 2 fold. Using novel single-molecule methods, we follow in real time and annotate the unfolding and refolding intermediates of Ross, a representative of this topology. This protein folds along a single reversible pathway involving the ordered and sequential organization of discrete and cooperative folding units or foldons. This strict order of events results from the formation of an initial, autonomously folding unit (primary foldon) followed by the subsequent organization of secondary foldons whose stabilities depend on their interactions with previously organized ones. This process, if generalized, represents a great simplification of the protein folding mechanism, as identification of foldons in nature and their folding hierarchy, should allow prediction of the folding pathway of any natural protein from its sequence.

## Introduction

Proteins play a central role in most cellular processes, from participating as structural units, to catalyzing and regulating most biomolecular reactions. To perform these functions, most proteins must fold into their native three-dimensional structures, and their ability to attain their correct folding is essential for maintaining the living state. In turn, protein misfolding can lead to aggregation that often result in amyloidogenic pathologies such as Parkinson’s or Alzheimer’s disease^1^. In 1961 Anfinsen showed that the folding of Ribonuclease A is dictated solely by its amino acid sequence and that the native state adopted by a polypeptide corresponds to its free-energy minimum^2^. In the following years, this understanding motivated a great effort to find the amino acid “code” that would make it possible to predict the structure of a protein from its amino acid sequence. This quest could be called the *strict* protein folding problem. However, seven decades of experimental and theoretical inquiry did not reveal simple rules to predict the structure from the sequence. This task has now been accomplished via AI and machine learning models trained on the large data set deposited in the protein data bank (PDB)^3,4^. However, the fundamental question, that can be called the *extended* protein folding problem, namely, how do proteins fold? remains unanswered^5–8^.

Based on theoretical considerations, it has been proposed that proteins attain their native state by diffusing over a potential energy surface often depicted in the form of a funnel; here the horizontal or x axis represents the accessible configurations of the polypeptide chain and the vertical or y axis the free energy of the molecule^7,9,10^. During folding, the chain starts at the top of the funnel at any given place and moves downward towards its bottom where the native state is located, decreasing simultaneously its configurational entropy and its free energy. In this representation, the molecule can in principle diffuse from top to bottom following any one of many possible folding paths, with the folding process being able to start anywhere along its sequence. Accordingly, the folding unit in this model is the amino acid residue, and any of them, or a combination of a few, can function as a nucleation site for the folding process. An alternative model has been proposed to rationalize the results of experiments using hydrogen–deuterium exchange with mass spectrometry (HDX-MS) of cytochrome C^5,11^. This study led these authors to propose that proteins do not start folding at any arbitrary place along their sequence, rather they attain their native state following one folding pathway with well-defined intermediates. Accordingly, these authors proposed that the folding of a protein is encoded in discrete folding units (sequences of 20 to 25 amino acids with different tendencies to fold autonomously, termed *foldons*^5,11^). In this model, the folding priorities (hierarchies) of foldons dictate the order in which they organize, thus defining a unique path of sequential folding intermediates.

A controversy has ensued around these opposite models^5–7^ and its resolution is of the utmost importance. However, because HDX-MS is a bulk or ensemble experiment, it does permit following individual folding trajectories of a single protein in real time that could provide the crucial discrimination between the two models. Although previous single molecule force spectroscopy experiments have revealed discrete folding and unfolding intermediates^12–16^, their identity could not be determined from their force-extension signature. Here, we present a novel single-molecule force spectroscopy approach using high-resolution optical tweezers to relieve this limitation and address the controversy.

## Results

### Folding through reversible intermediates

Our model is a *de novo*-designed protein PDB 2LV8, *Ross*^17,18^. Ross has the Rossmann2 × 2 fold which is composed of alternating four β-strands and four α-helices, β^1^α^2^β^3^α^4^β^5^α^6^β^7^α^8^ (**Fig. 1A, Table S1**). The fold is conserved, ancient, diverse and it is found in more than 20% of the proteins deposited in the PDB (examples in **Fig. S1**)^19–21^. Ross does not have cysteines, is stable^17^, and intermediates were previously detected in the optical tweezers^18^.

**Figure 1.**
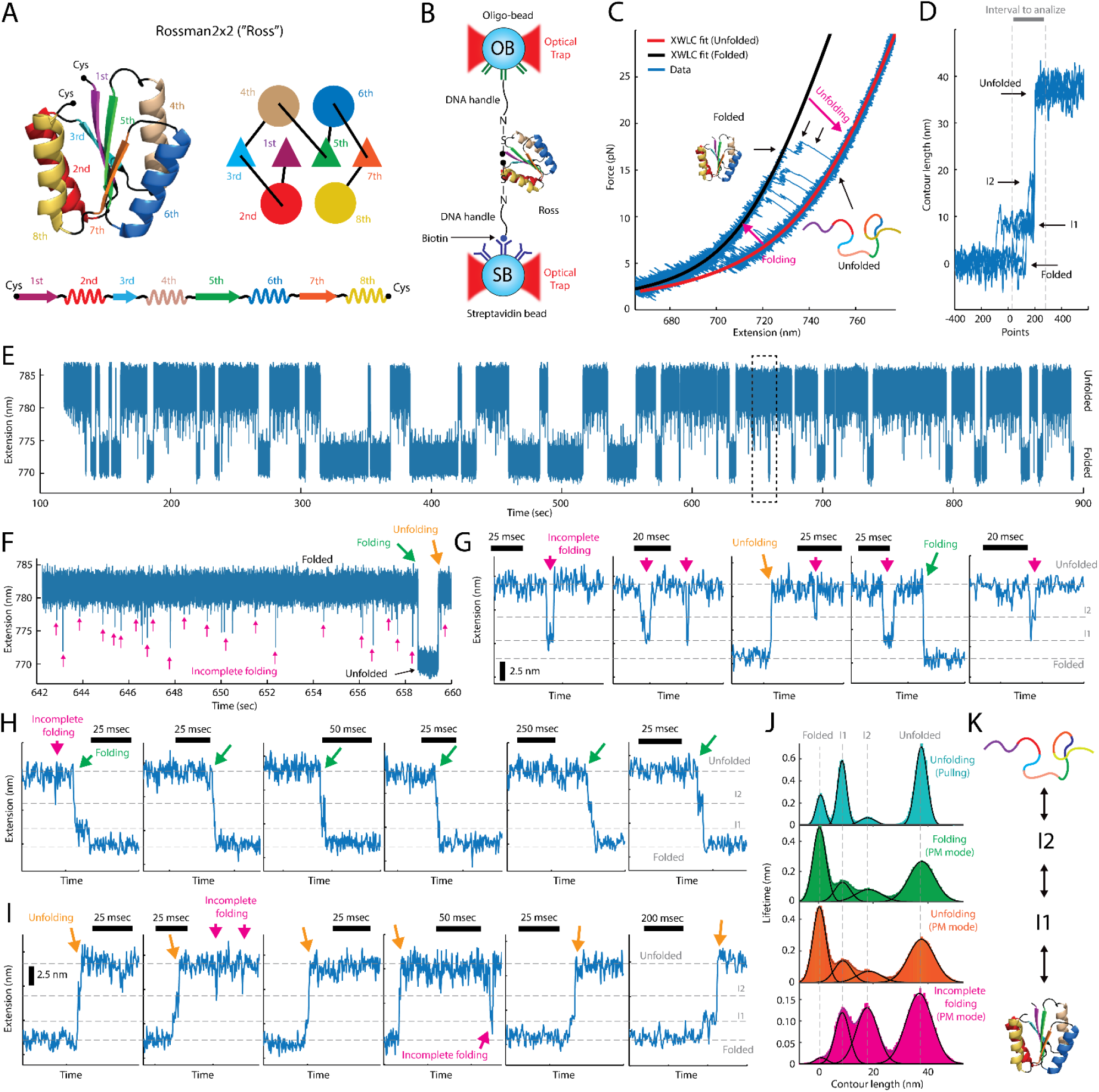
Discreate and defined intermediates during (un)folding of Ross. **A)** *Left*, Ross structure (PDB) with cys located at the C- and N-terminal. *Right*, simplified connectivity of secondary elements (triangles are β-strands and circles are α-helices) by loops (black lines). *Bottom*, linear connectivity of secondary elements. Each secondary structure is color-coded and numbered. **B)** Dual-trap Optical tweezers set-up to manipulate proteins (Ross in this case) via pulling/relaxing or passive mode (PM). Ross is tethered by two dsDNA molecular handles of ∼1000 bp via the covalent linkage of N-HS ester between the thiol group (cys) of the protein and the amino group of the 5’ end of the handle (see methods for details). Each dsDNA handle is then attached a polystyrene beads via a covalent bond and a biotin-streptavidin bridge. **C)** Five representative pulling/relaxing curves, extension (nm) vs force (pN), of Ross at the speed of ∼900 nm/s and the XWLC fitting to the data (blue, downsampled to 1250 Hz) when the protein is folded (black) and unfolded (red). **D)** Corresponding contour length Lc (nm) vs time (points) of the data (blue, downsampled to 2.5 kHz) from C after conversion using the XWLC. Folded, unfolded, and intermediates I1 and I2 are indicated with black arrows. Discontinuous vertical red line indicates the interval chosen for analysis of Lc histogram distribution. **E)** Representative PM data (blue, downsampled to 2.5 kHz), extension (nm) vs time (sec), of a single Ross. When Ross is unfolded, the tether has the larger extension (top); and when Ross is folded the tether experiences a compaction (bottom). **F)** Zoom in on the black dotted region from the trace in E. Small red arrows indicate incomplete folding attempts, whereas green and orange arrow indicated compete folding and unfolding. **G)** Five examples of incomplete folding (downsampled to 2.5 kHz) in Passive mode. Some events are accompanied by a complete folding or unfolding for reference of fully folded and fully unfolded states of Ross. Ross can fold into I2 or I1 (sometimes though I2) and fully unfold again via I2 or not. **H)** Six examples of folding of Ross in passive mode (downsampled to 2.5 kHz). **I)** Six examples of unfolding of Ross in passive mode (downsampled to 2.5 kHz). Scale in time and extension are indicated in G, H and I. **J)** Histogram of the Lc distribution of the unfolding of Ross during pulling (cyan), folding in PM mode (green), unfolding in PM mode (orange), and incomplete folding events during PM mode (red). See Table S2 for details. **K)** Model of (un)folding of Ross between the folded and unfolded states via two well defined intermediates I1 and I2. The color code is the same as in A.

We developed a high efficiency method to tether a single Ross molecule by its ends between a streptavidin and an oligo-modified bead, via dsDNA handles (**Fig. 1B, Fig. S2**, see methods). By moving the two beads repeatedly farther or closer from each other, mechanical force was applied to Ross, leading to unfolding/refolding of the protein through multiple pulling/relaxing cycles (**Fig. 1C, Fig. S3**). For each cycle we extracted the contour length (L_c_) of the protein resulting from the unfolding event (**Fig. 1D**, see methods). Analysis of the L_c_ histogram over n = 823 transitions showed two well-defined intermediates at ∼23% and ∼47% of the total L_c_ of Ross (∼37.6 nm), which we labelled as I1 and I2, respectively (**Fig. 1D, Table S2**). These intermediates are consistent with previous observations^18^.

Next, we sought to determine if the pathway defined by these intermediates is reversible. To this end, we performed passive mode experiments^22^ where the distance between the traps is held constant at a value in which Ross transits between its folded and unfolded states (**Fig. 1E**). Analysis of L_c_ around the folding and unfolding events (**Fig. 1F - G**) revealed two intermediates with the same L_c_ values observed in the pulling/relaxing experiments (**Fig. 1H, Table S2**). Therefore, the unfolding and refolding of Ross, either at equilibrium (passive mode) or out-of-equilibrium (pulling/relaxing), is a reversible process with cooperative and discrete intermediates I1 and I2. In sum, the unfolding of Ross is the reverse reaction of folding.

### Folding intermediates annotation and identification of foldons

To determine the identity of I1 and I2, we introduced a novel annotation method that involves generating seven variants each harboring an inserted segment of eighteen glycines (Gly-loop) between each two adjacent secondary structures of the protein. Variants are named Ross- i/i+1, when the Gly-loop is located between the i^th^ and the (i+1)^th^ secondary structure, e.g. Ross- 2/3 has the Gly-loop located between the 2^nd^ (α-helix) and the 3^rd^ (β-strand) secondary structures (**Fig. S4, Table S1)**. When a particular segment in the primary sequence of the protein containing the engineered Gly-loop is unfolded, the L_c_ of that transition should be lengthened by a distance equivalent to the L_c_ of the loop (18xGly ∼6.3 nm) relative to the same transition in the wild-type protein. Therefore, the position of the Gly-loop in that construct determines the part of the protein that unfolded during that transition (**Fig. S5, Fig. 2A**). Unlike the labelling of proteins with fluorophores, the unstructured Gly-loops have small effect on the mechanical stability of the protein (**Fig. 2B**) and its folding kinetics^23,24^, do not alter the structure of the protein as predicted by AlphaFold and Rossetta relaxation (**Fig. S4, S6**), and avoid the photophysical issues associated with fluorescence labels, such as photobleaching.

**Figure 2.**
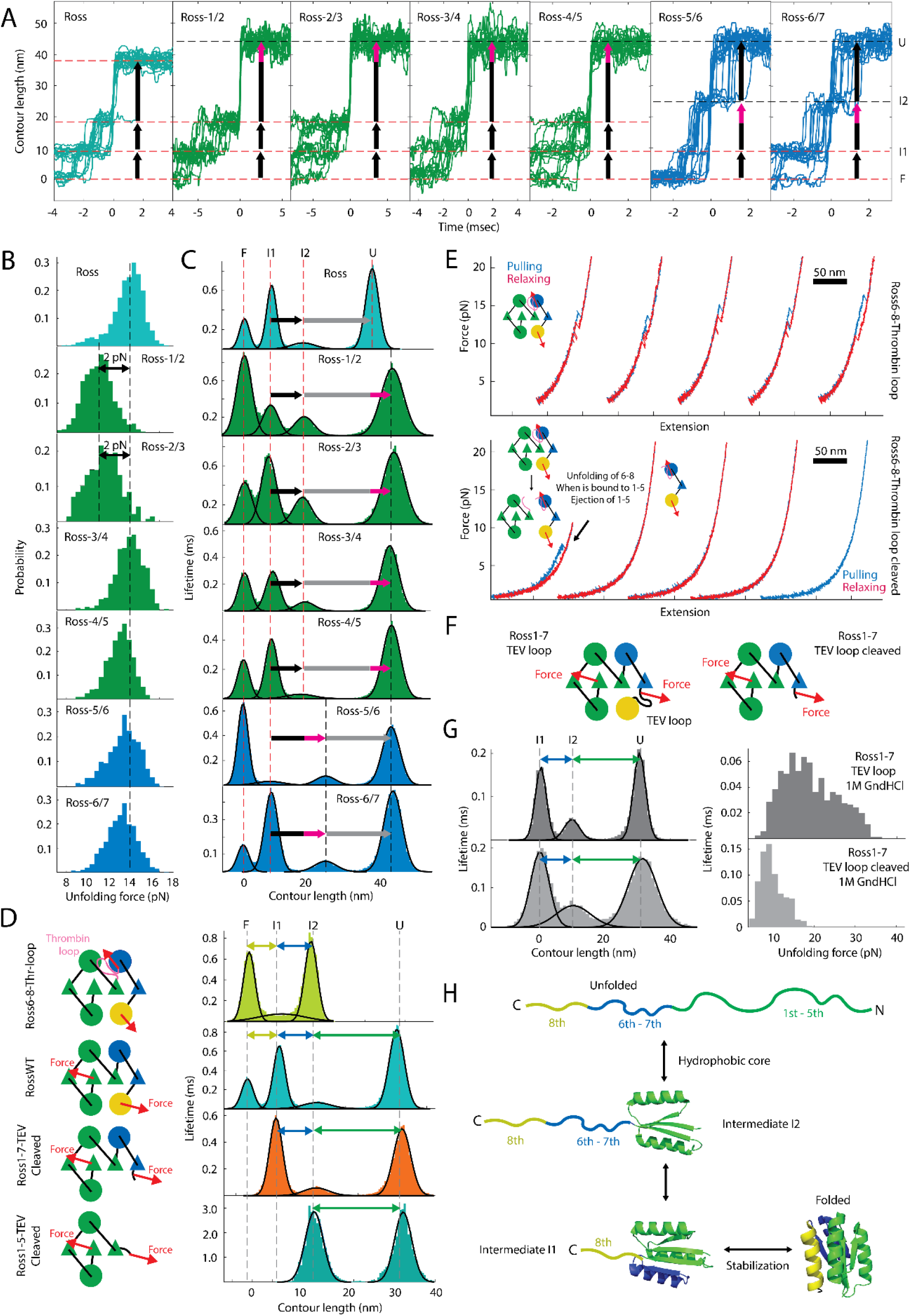
Dissection of the nature of the intermediates of the (un)folding of Ross. **A)** Twenty individual unfolding transitions (L vs time in nsec, downsampled to 6250 Hz from raw Data) of Ross and all the poly-Gly loop variants. Red discontinuous lines represent the mean Lc values of either folded (F), intermediates (I1 and I2), and unfolded (U) states. Black discontinuous lines represent the shifted Lc value of I2 and U from transitions I1 → I2 and I2 → U. **B)** Unfolding force distribution of all poly-Gly loop variants of Ross. Discontinuous line represents the mean value of unfolding force of Ross wild-type as reference. Only Ross-1/2 and Ross-2/3 show a significant difference in their mechanical unfolding. **C)** Histogram of Lc distribution of the unfolding transitions for all poly-Gly variants of Ross. Red lines represent the mean Lc values of either folded (F), intermediates (I1 and I2), and unfolded (U) states (see values in Table S3). Black arrows represent the change in contour length (ΔLc) between I1 to I2 not considering the poly-Gly loop. When the poly-Gly loop is also part of the unfolded region of Ross between I1 and I2, the lengthening of the ΔLc by 6.2 nm is depicted as the extension of the black arrow in purple color (this is also the case for the ΔLc between I2 and U, with the difference that when the poly-Gly is not part of the unfolded region of Ross, the arrow is depicted with grey color). **D)** *Left*, representation of constructs to study the first foldon alone (green, Ross1-5-TEV cleaved), the first and second (blue) foldons together without the third foldon (Ross1-7-TEV cleaved), and the second and third (yellow) foldons together in the presence of the first green foldon (Ross6-8-Thr- loop). *Right*, histogram of the Lc distribution of Ross6-8-Thr-loop (light green) has an intermediate that coincides with Lc value between the folded and I1 state of Ross (green). In the case of Ross1- 7-TEV cleaved, the folded, the intermediate, and unfolded states of the histogram of the Lc distribution (orange) coincide with the relative values of Lc of I1, I2 and folded states of Ross, respectively. Finally, there is no observable intermediate between the folded and unfolded state of Ross1-5-TEV cleaved (cyan), like the ΔLc between I2 and U of Ross. For all Lc histogram distributions, the color (yellow, blue and green) of the double arrow corresponds with the secondary structures of Ross unravel during a specific transition. **E)** *Top*, five consecutive pulling (blue) and relaxing (red) cycles, shifted by 100 nm each for clear visualization of the Ross6-8-Thr- loop construct. *Bottom*, Similar as top but the thrombin loop has been cleaved. In the first pulling, there is a rip indicating the 6^th^-8^th^ (β^6^α^7^β^8^) has a define and stable structure when it is interacting with first foldon (1^st^-5^th^ = β^1^α^2^β^3^α^4^β^5^). However, after the rip, the first foldon is ejected and therefore 6^th^-8^th^ does not fold after the first relaxing. 6^th^-8^th^ remains unfolded in subsequent pulling/relaxing cycles. **F)** Simplified diagram of the constructs containing the TEV cleavage sequence in a loop located between 7^th^/8^th^ of Ross: uncleaved (left) and cleaved (right). **G)** *Left*, histogram of Lc distribution of the unfolding transitions for the uncleaved (top) and cleaved (bottom) construct, both in normal buffer conditions with 1 M guanidinium chloride. Discontinuous lines represent the F, unfolded and intermediate. *Right*, unfolding force distribution of the corresponding conditions in F. **H)** Folding mechanism of Ross.

The pulling/relaxing experiments of all Ross Gly-loop variants (**Fig. S7, Fig. 2B**) showed that ΔL_c_ of the transition between I2 and U is extended by ∼6.4 nm in the case of Ross-1/2, -2/3, - 3/4, and -4/5, indicating that the first folding unit of Ross involves the cooperative organization of β^1^α^2^β^3^α^4^β^5^ (1^st^ foldon). Next, ΔL_c_ between I1 and I2 is extended by ∼6.4 nm in Ross-5/6, and -6/7, indicating that the second folding unit involves the cooperative grouping of α^6^β^7^ (2^nd^ foldon, **Fig. 2C, Table S3**). Finally, it is expected that Ross-7/8 would show that ΔL_c_ between F and I1 is extended in the presence of the Gly-loop. However, Ross-7/8 did not express well. Changing the loop sequence, induction conditions, inductor, and induction strain did not help (data not shown). These results suggest that construct Ross-7/8 is not stable, even though the 8^th^ secondary structure (α^8^) must necessarily be the last cooperative folding step of Ross (3^rd^ foldon). To confirm this inference, we performed molecular dynamics simulations of the pulling of Ross (see methods). These simulations confirm that β^1^α^2^β^3^α^4^β^5^ is the first foldon whereas α^6^β^7^ and α^8^ are the second and the third foldons respectively (**Fig. S8**). Simulations also show that the Gly-loops do not alter the unfolding pathway of Ross (**Fig. S8**). All experiments above were carried out at 300 mM KCl. We observed the same folding units when the experiments were performed at 25 mM KCl or at 300 mM KCl plus 1 M Guanidinium, although in this latter condition the unfolding and refolding forces were noticeably reduced (**Fig. S8, Table S3**). Overall, the folding of Ross involves a single well-defined pathway punctuated by discrete intermediates that sequentially and cooperatively incorporate different groups of secondary structures, and that are visited in strict opposite order during unfolding.

### Coupling between foldons defines their folding order

One way to rationalize the strict folding order observed for Ross is the existence of a hierarchy of autonomy among its foldons. Accordingly, folding always starts with an initial foldon, which can adopt its structure autonomously, followed by organization of units whose stability depend on their interaction and energetic coupling with previously organized ones. To test this idea, we repeated the unfolding/relaxing experiments of the 1^st^ foldon (β^1^α^2^β^3^α^4^β^5^), the 1^st^ and the 2^nd^ foldons together (β^1^α^2^β^3^α^4^β^5^-α^6^β^7^), and the 2^nd^ and 3^rd^ foldons together (α^6^β^7^-α^8^), either isolated or when still covalently linked to their respective complementary regions, to determine if they can fold autonomously in either condition (**Fig. S10, S11, Fig. 2D**, see methods). To this end, we developed a method to isolate fragments of Ross *in-situ* in the optical tweezers. We engineered (i) two cysteines surrounding the desired region in the full sequence of the protein, and (ii) a protease loop sequence between the desired and complementary sections of Ross. We produce the protein as a full-length product, cleave the loop with the protease, and attach dsDNA handles to the cysteines. In the optical tweezers, the first pulling unfolds the desired section of Ross and breaks its non-covalent interactions with the complementary region of the protein. As a result, the latter is released to the solution, leaving the desired region of Ross isolated *in-situ* to be manipulated in subsequent relaxing/pulling cycles.

Experiments showed that the 1^st^ foldon, once unfolded, can refold into a stable structure in the presence or in the absence of the 2^nd^ and 3^rd^ foldons (**Fig. S12A**). Similarly, the 1^st^ and 2^nd^ foldons always fold in the presence or absence of the 3^rd^ foldon (**Fig. S12A**). However, the 2^nd^ and 3^rd^ foldons by themselves (when not linked to the 1^st^ foldon) do not reveal an unfolding or refolding transition (**Fig. 2E**). Furthermore, pulling the 2^nd^ and 3^rd^ foldons while connected to β^5^ (β^5^α^6^β^7^α^8^) does not fold (**Fig. S13**).

The above results indicate that the 1^st^ foldon can organize autonomously and as such it initiates the folding of Ross (we designate it as the *primary* foldon). The primary foldon is required for the folding of the 2^nd^ foldon, while the organization of the first two foldons is required for the folding of the 3^rd^. We designated the last two foldons as *secondary* i.e., as foldons that are thermodynamically unstable unless a previous foldon of higher priority is already organized. Consistent with the energetic coupling hypothesis, analysis of the pulling/relaxing data (**Fig. S14**) shows that the unfolding force of the combined 1^st^ and 2^nd^ foldons is significantly higher in the presence of the 3^rd^ foldon than in its absence (**Fig. 2F-G, Fig. S15**). Importantly, the L_c_ obtained from pulling the isolated 1^st^ foldon, the 1^st^ and 2^nd^ foldons together, and the 2^nd^ and the 3^rd^ foldons (linked to the 1^st^) coincided with the ΔL_c_ values of the folding units derived from the poly-Gly loop experiments (**Fig. 2C, Table S3, Fig. S16**). In sum, the cooperative folding units of Ross display different degrees of folding autonomy which translates into a well-defined folding order (**Fig. 2H**). The primary foldon adopts its structure by itself because its amino acid sequence encodes the necessary intra-foldon interactions that permit it to fold autonomously, cooperatively and stably. In contrast, the structural organization of secondary foldons depends on inter-foldon interactions with previously organized foldons, and they are not thermodynamically stable by themselves. In general, folding of successive units provide increasing stabilization for those organized previously. This energetic coupling leads to the ordered, unique, and hierarchical folding path observed for this protein.

### Folding intermediates formed at near zero force

Applying tension on the protein from the ends may bias its folding pathway, preventing for example an early event involving the interaction of the N- and C-terminus. To rule out this scenario, we developed the Rapid Force Quench (RFQ) method. Briefly, starting with the folded tethered protein, we rapidly increase the trap separation to a fixed value (**Fig. 3A**), raising the tension applied to the protein, leading to its unfolding. Next, we suddenly drop (quench) the force to near zero pN (**Fig. S17**) for a short period of time to allow for the folding of the protein under these conditions, before increasing again the force (**Fig. 3A left, inset**). The time the protein experiences near zero force (the quench time) must be of the order of the time required for the protein to fold, to capture the protein in a partially folded structure on its way to the native state. The sudden increase of the force after the quench time prevents the folding from continuing and is used to determine the nature of the intermediate formed under zero force from the size of its contour length (**Fig. 3A right, inset**). We performed these experiments in a time-shared, dual-trap high-resolution optical tweezers^25,26^, which affords a quick motion of the optical trap (∼10 µsec), which determines the time resolution of the RFQ method.

**Figure 3.**
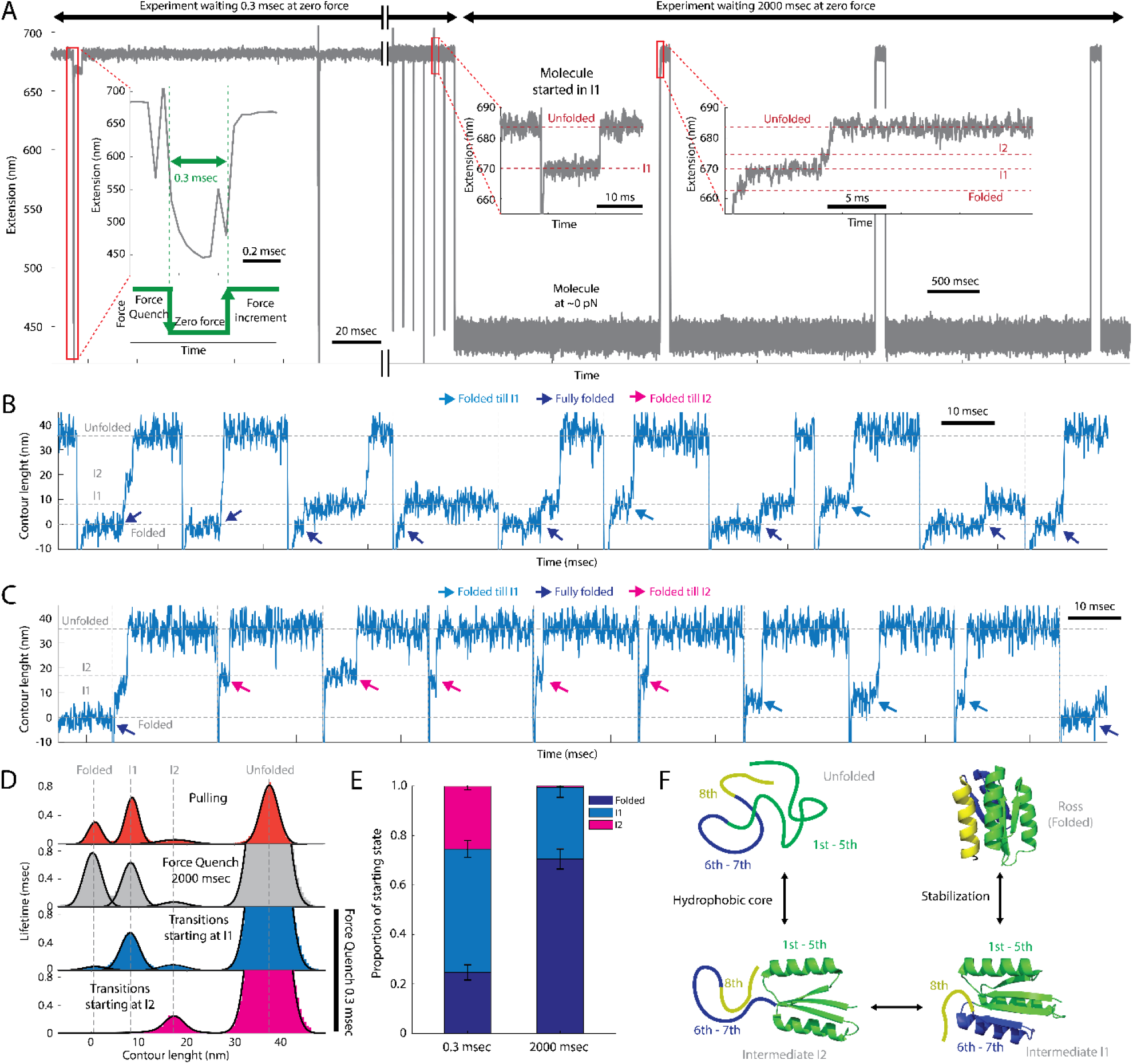
Folding intermediates at near zero force. **A)** Representative trace of extension (nm) vs time (msec) of one Ross molecule waiting 0.3 msec (experiment) and later 2000 msec (control) at near zero force. *Left*, expanded section of two representative quenching events. *Inset*, zoom in to show that in one quenching time, the molecule was around 0.3 msec at a tether extension below ∼550 nm where the force experienced by the molecule is near zero (**Fig. S17**). *Right*, same trace showing when the last cycles using a quenching time of 0.3 msec and, for the same molecule, we use rather 2000 msec at ∼0 pN as a control experiment. *Zoom in*, of unfolding transitions, if any, after waiting 0.3 msec or 2000 msec. For the former, Ross folded till I1 and for the latter, Ross fully organized because it went through I1 and I2 upon unfolding (Intermediates, unfolded and folded states are indicated with horizontal discontinuous red lines). **B)** Contour length (nm) vs time (msec) of continuous unfolding transitions of partially organized sub-structures of Ross at near zero force for 0.3 msec of waiting time. **C)** Contour length (nm) vs time (msec) of continuous unfolding transitions of partially organized sub-structures of Ross at near zero force for 2000 msec of waiting time. In both case C and D, the data is at 25 kHz, discontinuous vertical gray lines are the separation between cycles, and discontinuous horizontal gray lines are the reference of folded, intermediates (I1 and I2) and unfolded states of Ross. **D)** Contour length histogram distributions of Ross from the analysis of pulling (orange), all transitions of RFQ at 2000 msec (gray), transitions starting at either I1 (blue) or I2 (purple) of RFQ at 0.3 msec. Fitting to Gaussian is shown is black lines for each histogram. Discontinuous vertical gray lines are to show all the respective states of Ross during unfolding in all the histogram are similar. **E)** Stacked bar chart of the probability of finding Ross to be fully folded (blue) or partially organized in I1 (blue) or I2 (purple). Error bar is the SEM calculated as the 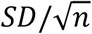 from three independent Ross molecules (n = 3). **F)** Folding mechanism of Ross at near zero force.

For Ross, the folding time is ∼1.5 msec and the time between intermediates is on the order of hundreds of µsec^18^. Therefore, we used quench times of 300 µsec, followed by application of ∼12 pN for 100 msec to unfold and analyze the parts of the protein folded during quench time (**Fig. 3B**). This quenching cycle was repeated several times. Each tethered molecule was also subjected to a control experiment in which the protein is allowed to fold under zero tension for 2 sec, after which it is pulled to confirm the presence of intermediates I1 and I2 as a reference (**Fig. 3B**). By visual inspection, we find that the molecules subjected to RFQ display significantly shorter unfolding transitions in length than that of the fully folded protein (as observed in the control experiment, **Fig. 3C-D**). Analysis of the Lc distribution of those states reveal that they correspond to intermediate I1 or I2 (**Fig. 3E, S18, Table S4**). From all the transitions observed, ∼25% started in I2, ∼50% in I1 and ∼25% correspond to the fully folded protein. In the control experiment, ∼30% started in I1, ∼70% in the fully folded state and we did not observe transitions starting at I2 (**Fig. 3F**).

In summary, by implementing the RFQ method, we were able to capture Ross midway during its path to the fully folded state under zero tension. Furthermore, the first intermediate formed at near zero force coincides with I2. We conclude that the Ross folds via the organization of β^1^α^2^β^3^α^4^β^5^ (I2) and transits through I1 before attaining its native state at zero tension (**Fig. 3G**), and that force as denaturant does not bias its folding path.

### Identification of a hidden foldon

Molecular dynamics pulling simulations of Ross and its poly-Gly loop variants suggest that β^1^α^2^β^3^α^4^β^5^ unfolds via the transient intermediate β^1^α^2^β^3^ that last from one to very few frames (**Fig. S8**), which is presumably too short-lived to be detected in our experiments. To determine if β^1^α^2^β^3^ can indeed fold as an independent unit, we used the protease loop single-molecule assay described above to isolate β^1^α^2^β^3^ *in-situ* in the optical tweezers (**Fig. 4A-C**). In the first pulling, β^1^α^2^β^3^ (bound to α^4^β^5^α^6^β^7^α^8^) displayed a cooperative unfolding transition at forces 15 – 37 pN, like the uncleaved construct (**Fig. 4D**). This force is required to unfold β^1^α^2^β^3^ and to break its non-covalent interactions with α^4^β^5^α^6^β^7^α^8^ (**Fig. 4E**). After the release of the latter (**Fig. 4E**), many of the subsequent pulling/relaxing curves looked like bare DNA (**Fig. 4C**). However, a subset of the relaxing curves of β^1^α^2^β^3^ isolated, showed refolding transitions in the range of 4 – 7 pN, bringing them to the first pulling curve of β^1^α^2^β^3^ when it was still bound to α^4^β^5^α^6^β^7^α^8^ (**Fig. 4F**). Consistently, pulling of β^1^α^2^β^3^ reveal unfolding transitions in the same range of forces (**Fig. S19**). Lastly, pulling simulations of β^1^α^2^β^3^ in isolation also show it to be resilient under force (**Fig. S20**).

**Figure 4.**
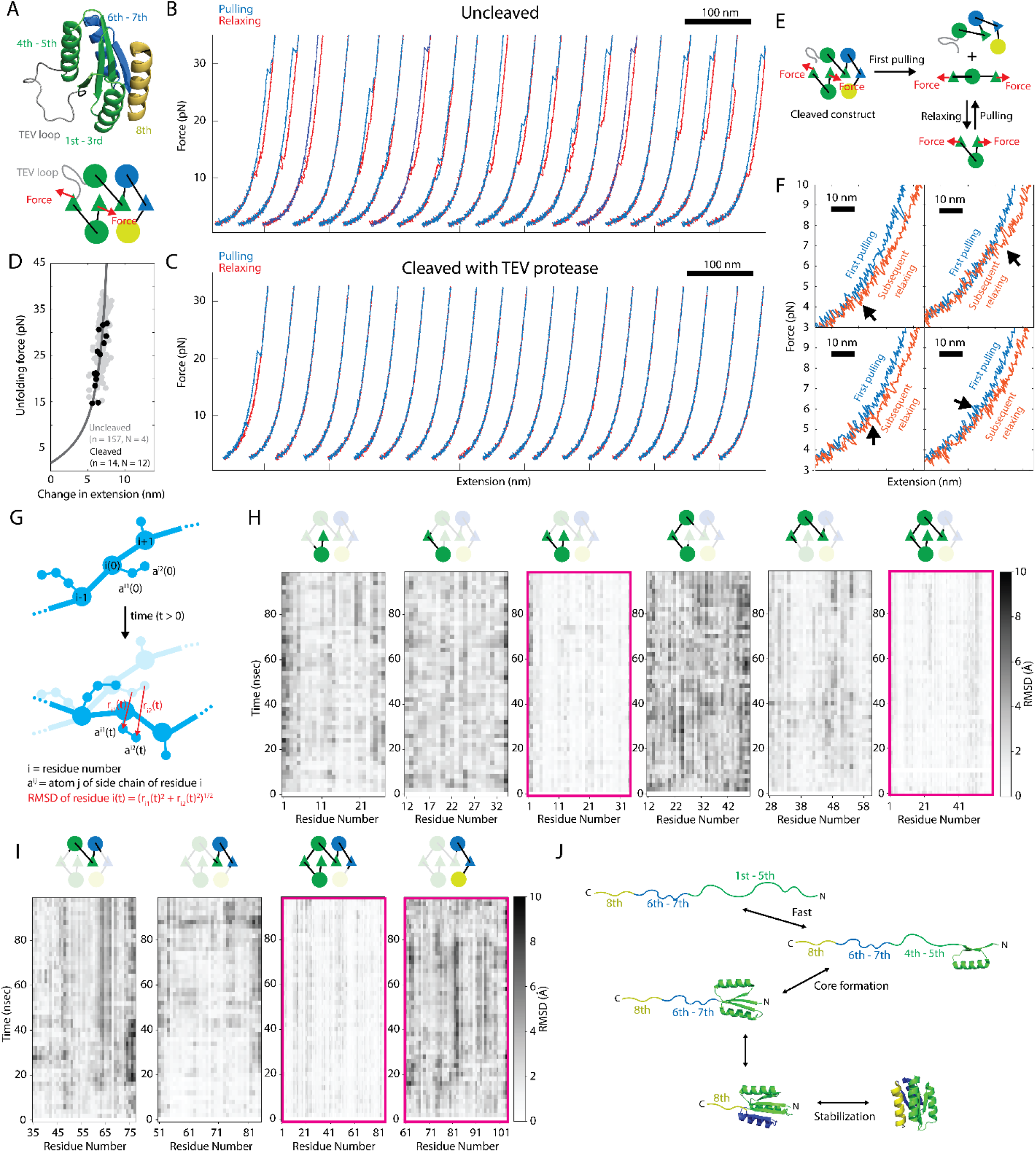
Folding of Ross starts with the organization of the three secondary structures in the N-terminus. **A)** Alpha-fold prediction (*top*) and diagram (*bottom*) of the Ross1-3TevLoop construct sequence. Color code corresponds to the folding units found using the poly-Gly loop method: green for β^1^α^2^β^3^α^4^β^5^, blue for β^6^α^7^ and yellow for α^8^. **B)** Representative 20 consecutive pulling (cyan) and relaxing (red) cycles, at 200 nm/s between 1.5 to 35 pN with 1 sec waiting time at low force, for Ross1-3TevLoop uncleaved. **C)** Representative 20 consecutive pulling (cyan) and relaxing (red) cycles from the first pulling for Ross1-3TevLoop cleaved with TEV protease (same conditions as B). **D)** WLC analysis of the change in extension vs unfolding force of all the transitions observed (N = 4 molecules displaying 157 transitions in total) for Ross1-3TevLoop uncleaved (gray). The single unfolding transition (black) observed in the first pulling (N = 12 molecules and 14 rips) for Ross1-3TevLoop cleaved. **E)** Scheme of the events happening during the only rip observed in the cleaved construct. In the first pulling, there is a rip indicating the 1^st^- 3^rd^ (β^1^α^2^β^3^) is a highly stable structure when it is interacting with its complementary region (4^th^- 8^th^ = α^4^β^5^α^6^β^7^α^8^). However, after the rip, α^4^β^5^α^6^β^7^α^8^ is ejected and therefore, in subsequent relaxing and pulling cycles, β^1^α^2^β^3^ is isolated. **F)** Four examples of clear folding transitions (downsampled to 50 Hz) of β^1^α^2^β^3^ during relaxing (red) compared to the first pulling (blue). **G)** Simplified diagram of calculation of the RMSD value of each residue at a given time (nm) vs the start of the equilibrium simulation (time 0). **H)** Variation of RMSD from 0 to 100 ns of all aminoacids of the construct tested in simulations related to 1^st^-5^th^ (β^1^α^2^β^3^α^4^β^5^): β^1^α^2^, α^2^β^3^, β^1^α^2^β^3^, α^2^β^3^α^4^, β^3^α^4^β^5^ and β^1^α^2^β^3^α^4^β^5^. **I)** Variation of RMSD from 0 to 100 ns of all aminoacids of the construct tested in simulations related to structed outside the folding unit 1^st^-5^th^: α^4^β^5^α^6^, β^5^α^6^β^7^, α^6^β^7^α^8^, and β^1^α^2^β^3^α^4^β^5^α^6^β^7^. **J)** Complete folding mechanism of Ross.

To explore whether β^1^α^2^β^3^ can fold by itself in solution, we developed and performed folded stability molecular dynamics simulations. We created truncations of Ross and determined whether they unfold quickly or remain folded in explicit solvent for the 100 nsec of the simulation (**Fig. 4G, Fig. S21**, see methods). We found that β^1^α^2^ and α^2^β^3^ unfold in less than 10 nsec, similar to α^6^β^7^α^8^ (**Fig. 4H-I**), which is known to be unstructured in isolation (**Fig. 2D**). However, β^1^α^2^β^3^, like β^1^α^2^β^3^α^4^β^5^ (“I2”) and β^1^α^2^β^3^α^4^β^5^α^6^β^7^ (“I1”), which display cooperative (un)folding transitions in the optical tweezers (**Fig. 2C**), remained stable beyond the 100 nsec of the simulation (**Fig. 4H- I**). Furthermore, other triplet combinations (α^2^β^3^α^4^, β^3^α^4^β^5^, α^4^β^5^α^6^, β^5^α^6^β^7^), were also unstable (**Fig. 4H-I**). Altogether, these results indicate that β^1^α^2^β^3^ can fold autonomously in solution and under force.

In summary, Ross folds via a cooperative, unique, and hierarchical sequence of events, whose order is strictly reversed during unfolding. The complete folding mechanism of Ross is then (**Fig. 4J**):

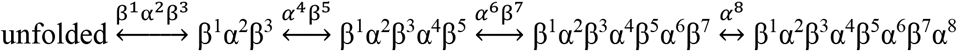

## Discussion and conclusions

The results presented here show that Ross, a member of Rossmann2 × 2 family and present in nearly one fifth of the proteins in the PDB, folds following a single path characterized by the formation in strict order of several discrete intermediates. This result supports the Foldon model and shows that the observed folding order of this protein arises from the distinct folding hierarchy of its constituent foldons.

The existence of foldons can explain why the nearly 200,000 structures currently in the PDB consistently contain a small number of repeated motifs (αα, β-hairpin, βαβ, etc). Their presence suggests that short runs of different amino acid sequences have a natural tendency to adopt similar spatial organizations and to function as folding units with some degree of autonomy. It is tempting to speculate that by combining these short sequences with different stabilities throughout evolution, larger and more complex proteins, possessing the ability to fold in a modular fashion, eventually emerged. This modular architecture could have greatly simplified the folding process of proteins and their search for the native structure. Interestingly, it is likely that such modularity eventually extended through different scales of folding. Larger proteins are seen to segregate into multiple globular domains that attain their native form in a well-defined order, often requiring some degree of inter-domain coupling^15,16^. Again, segregation of large proteins into distinct globular folding domains likely simplifies their folding, with these larger cooperative folding domains functioning as *superfoldons*.

Future studies of other conserved folds (e.g. Ferredoxin, SH3, β-sandwich, β-grasp) using the approach developed here, in combination with HDX-MS, will provide a comprehensive understanding of the general rules of protein folding. Validation of the foldon model for other conserved folds would represent a great simplification of the protein folding mechanism. In this scenario, folding proteins would not be described as polymers of aminoacid but as a oligomers of foldons. The eventual identification and classification of new foldons, their relative folding hierarchy, and their interdependence with other foldons would make it possible, in principle, to predict the folding path of any given natural protein from inspection of its sequence alone.

## ACKNOWLEDGEMENTS

We thank the assistance of Rohit Satija, Vijay Prathigudupu, Dilara Zenel, Alicia Li, and Benjamin Kuznets-Speck during various stages of biochemistry, cloning of proteins, data collection and analysis. We thank the valuable discussion with other members of the Bustamante laboratory. This research was supported by the National Institutes of Health grants R01GM071552 and R01GM032543.

## CONTRIBUTIONS

**RS:** Guided the project, conceived the experiments, developed the biochemical methods, perform the biochemistry, developed the single-molecule methods, collected and analyzed the optical tweezers data, wrote the paper. **JK:** performed biochemistry, collected and analyzed optical tweezers data. **AT:** Developed the instrumentation, developed the software for optical tweezers data collection and analysis, analyzed the data. **SK**: developed and performed the simulations. **CV:** performed biochemistry and collected the optical tweezers data. **BH:** performed the biochemistry and collected optical tweezed data. **DP**: Performed Rosetta relaxations. **CB:** Conceived the idea, guided the project and wrote the paper.

## Notes

### Competing Interest Statement

The authors have declared no competing interest.

### Summary of Updates

The manuscript has been revised manuscript and the supplemental files have been updated.

