## Supplementary Material for "The Rossmann2 × 2 Fold Attains its Native Structure Via a Defined Pathway of Sequential and Cooperative Folding Units"

**Fig. S1:** Some examples of protein and complexes in cartoon representation found the PDB that contain the Rossmann2x2 fold within its topology. The PDB code and the name of the protein are indicated above each structure. The color code corresponds to the same in Ross. Extra secondary structures to the Rossmann2x2 in each protein is depicted in grey color.

**Fig. S2:** Simplified scheme of the method of labelling proteins using the dsOligo with the N-hydroxysuccinimide (NHS) ester (GMBS). See methods for details.

**Fig. S3:** Representative example of four consecutive pulling (blue) and relaxing (orange) curves of Ross between 1.5 and 30 pN. Raw data collected at 25 kHz is shown in grey color. Downsampled data at 250 Hz is shown in the color mentioned above. *Inset.* Zoom in of two unfolding transitions depicting one of two clear intermediates, labelled as I1 and I2.

**Fig. S4:** Alpha fold prediction in cartoon representation of the poly-Gly loop variants of Ross. Color code is the same as Ross. As shown in bottom right, all predicted structures align well with Ross and among themselves.

**Fig. S5:** Explanation of how the poly-Gly loop method can annotate and determine the secondary structure components of a particular intermediate. *Left.* A, B and C represent three α-helices, shown in different colors, which are connected by curved black lines (turns). The poly-Gly loop is depicted in a discontinuous red lines. *Medium*. Scheme of force vs extension curves (gray color) depicting one clear intermediate I. If the poly-Gly loop is located between helix B/C and the second transition is lengthened, it means the folding mechanism is unfolded ⇄ BC ⇄ ABC (*right*). In the complementary experiment. If the poly-Gly loop is located between helix A/B and first transition is lengthened, it means the folding mechanism above still holds (*right*). As a result, in the absence of the poly-Gly loops, the mechanism of the protein is unfolded ⇄ BC ⇄ ABC (*right top*).

**Fig. S6:** Rosetta relaxation (cartoon representation) of the Alpha fold predictions of the poly-Gly variants of Ross. The color code is shown as in Ross.

**Fig. S7:** Fifteen consecutive pulling (blue) and relaxing (red) cycles of representative molecule per poly-Gly loop variants of Ross. All plots are in force (pN) vs extension (nm). Scale bar is shown on the top left. Each cycle of pulling/relaxing is shifted 100 nm to the left for visualization. Data has been downsampled to 250 Hz.

**Fig. S8:** Stable intermediates observed during pulling simulations for Ross and its poly-Gly loop variants (from left to right) from the ends. The color code is shown as the first five secondary structures in green (foldon 1), the $6^{th}$ and $7^{th}$ are in blue (foldon 2) and the last helix in yellow (foldon 3). In all trajectories, we observed a very short-lived intermediate composed of the first three secondary structures. The poly-Gly loop does not affect the unfolding mechanism.

**Fig. S9:** Histogram of the unfolding force (pN) and Lc (nm) distribution of unfolding transitions for all poly-Gly variants of Ross at 25 mM KCl (*left*) and 300 mM KCl with 1 M of Guanidinium (*right*). Red lines represent the mean Lc values of either folded (F), intermediates (I1 and I2), and unfolded (U) states (see values in Table S3). Black arrows represent the change in contour length (ΔLc) between I1 to I2 not considering the poly-Gly loop. When the poly-Gly loop is also part of the unfolded region of Ross between I1 and I2, the lengthening of the ΔLc by 6.2 nm is depicted as the extension of the black arrow in purple color (this is also the case for the ΔLc between I2 and U, with the difference that when the poly-Gly is not part of the unfolded region of Ross, the arrow is depicted with grey color).

**Fig. S10:** Alpha fold prediction in cartoon representation of the variants of Ross containing the Thrombin or TEV cleavage sequence in a loop. Color code is the same as Ross. As shown in bottom right, all predicted structures align well with Ross and among themselves.

**Fig. S11:** Rosetta relaxation (cartoon representation) of the Alpha fold predictions of the variants of Ross containing the Thrombin or TEV cleavage sequence in a loop. The color code is shown as in Ross.

**Fig. S12:** Fifteen consecutive pulling (blue) and relaxing (red) cycles of a representative molecule of the first five (*top*) and seven (*bottom*) secondary structures of Ross isolated. All plots are in force (pN) vs extension (nm). Scale bar is shown on the top left. Each cycle of pulling/relaxing is shifted 100 nm to the left for visualization. Data has been downsampled to 250 Hz.

**Fig. S13: A)** Schematic representation of constructs to study Ross pulling between the 5$^{th}$ and 8$^{th}$ secondary structures (Ross5-8) in presence (*top*) and in the absence (*bottom*) of the first four secondary structures of Ross. B) Five consecutive pulling (blue) and relaxing (red) cycles of a representative molecule of the constructs shown in A. All plots are in force (pN) vs extension (nm). Scale bar is shown on the top right. Each cycle of pulling/relaxing is shifted 100 nm to the left for visualization. Data has been downsampled to 250 Hz.

**Fig. S14:** Fifteen consecutive pulling (blue) and relaxing (red) cycles of a representative molecule of Ross being pulled between the 1$^{st}$ and 7$^{th}$ secondary structures (*top*), when there is a TEV cleavage sequence between 7$^{th}$ and 8$^{th}$ secondary structures (*middle*) and in the absence of the 8$^{th}$ secondary structure (*bottom*). All plots are in force (pN) vs extension (nm). Scale bar is shown on the top left. Each cycle of pulling/relaxing is shifted 100 nm to the left for visualization. Data has been downsampled to 250 Hz.

**Fig. S15: A**) Schematic representation of constructs to study Ross in the absence of the 8$^{th}$ secondary structure, Ross1-7, and pulling between 1$^{st}$ and 7$^{th}$ secondary structures. *Left*, control construct when the 8$^{th}$ secondary structure is present. *Middle*, when 8$^{th}$ is present but there is a TEV cleavage sequence between 7$^{th}$ and 9$^{th}$ secondary structures. *Right,* Ross construct missing 8$^{th}$. **B)** Unfolding force (pN) distribution of each construct shown in A. **C)** Lc distribution of unfolding events of each construct shown in A. Experiments were carried out in the HMK buffer and in the presence of 1 M Guanidinium.

**Fig. S16:** Stable intermediates observed during pulling simulations for Ross, and Ross in the absence of $6^{th}$-$7^{th}$-$8^{th}$ and in the absence of $8^{th}$ secondary structures, Ross1-5 and Ross1-7 respectively.

**Fig. S17:** Force (pN) vs extension (nm) plot of the initial pulling (black) and relaxing (orange) of Ross at 100 nm/s to determine trap positions to perform the RFQ experiments. Data downsampled to 625 Hz. Discontinuous vertical lines in grey color are the trap positions to have the tether extended ~670 nm for unfolding of partially folded substructures at ~12 pN and at ~450 nm for near zero to allow folding. *Top*, simplified energy diagram of folding of Ross (green) with I1 and I2 intermediates indicated. Once tether distances are defined, we move the laser quickly between those two positions (purple arrow for extension, cyan arrow for quenching).

**Fig. S18:** Contour length (nm) vs time (msec) of continuous unfolding transitions of partially organized sub-structures of Ross at near zero force for 2000 msec and 0.3 msec of waiting time. Each type of transition is specified in each plot.

**Fig. S19:** Twelves examples of clear unfolding transitions (downsampled to 50 Hz) of the first three secondary structures of Ross (Ross1-3) during subsequent pulling (blue) compared to the first pulling (black). Each plot is in force (pN) vs extension (nm). The scale bar in all cases is 10 nm.

**Fig. S20:** Pulling simulation of the first three secondary structures of Ross (Ross1-3). *Left.* Force (pN) vs extension (nm) curve of a single run. *Right.* Corresponding structures of the construct under different forces before and after the unfolding transition.

**Fig. S21:** Variation of RMSD from 0 to 100 ns of all aminoacids of the constructs (above each kymograph) tested in two more equilibrium stability simulations (A and B) in addition to the runs presented in Fig. 3H-I.

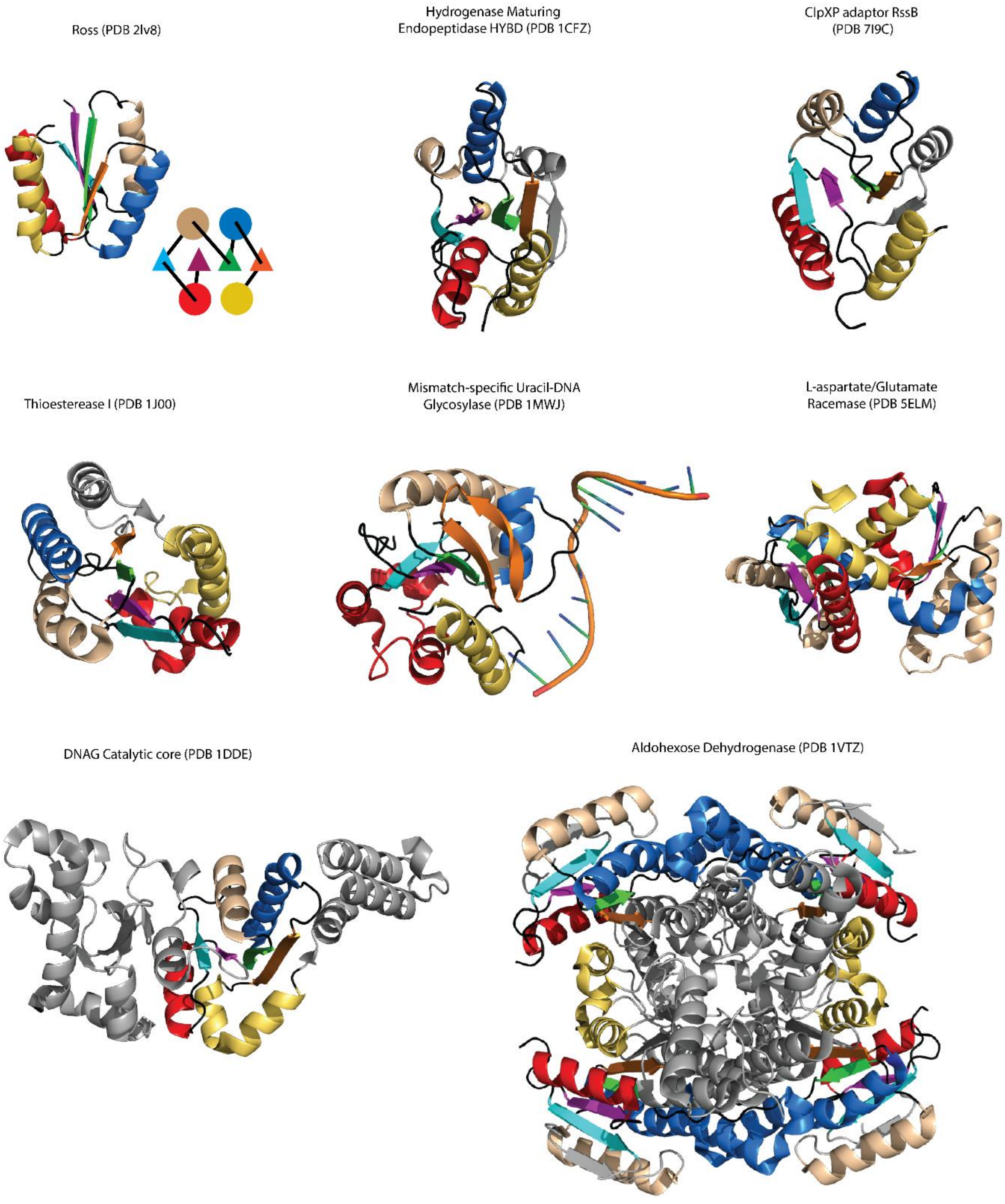
Ross (PDB 2lv8)
Hydrogenase Maturing
Endopeptidase HYBD (PDB 1CFZ)
ClpXP adaptor RssB
(PDB 7I9C)
Thioesterease I (PDB 1J00)
Mismatch-specific Uracil-DNA
Glycosylase (PDB 1MWJ)
L-aspartate/Glutamate
Racemase (PDB 5ELM)
DNAG Catalytic core (PDB 1DDE)
Aldohexose Dehydrogenase (PDB 1VTZ)

**Fig. S1**

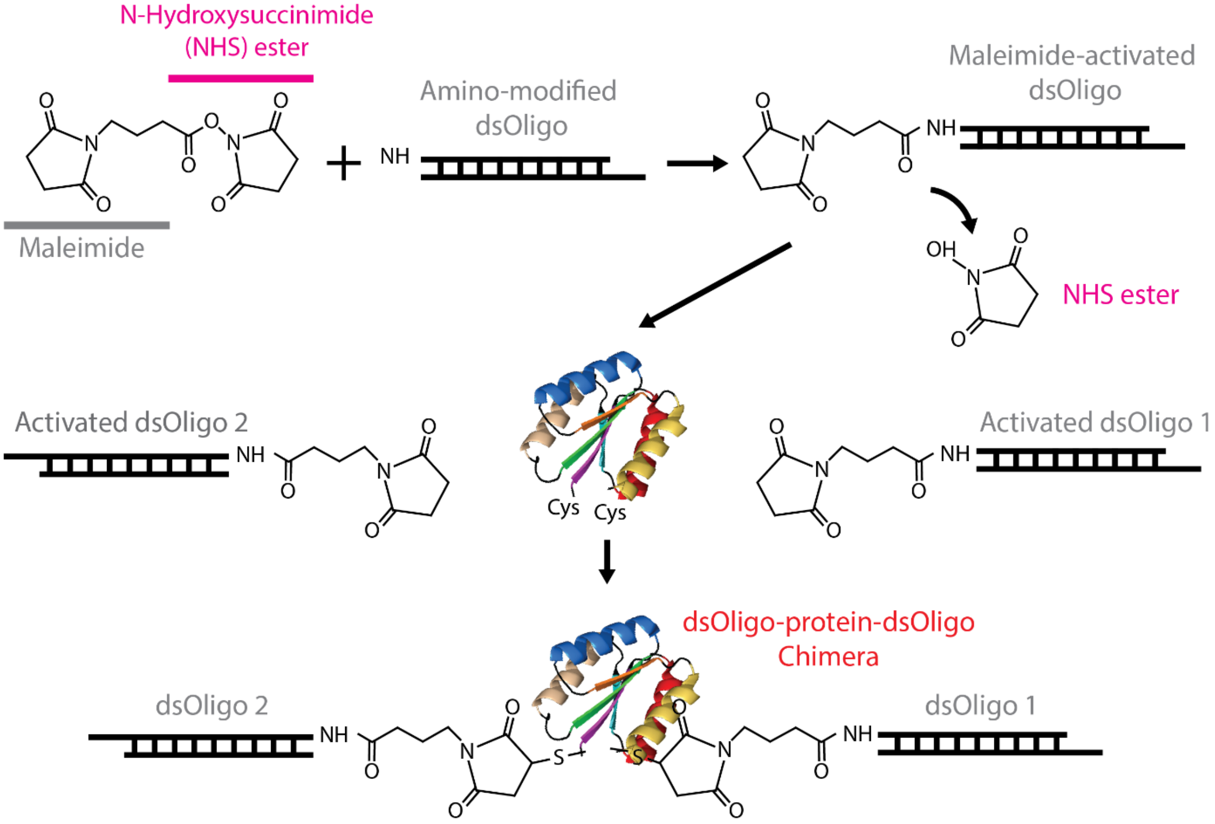
N-Hydroxysuccinimide
(NHS) ester
Maleimide
Amino-modified
dsOligo
Maleimide-activated
dsOligo
NHS ester
Activated dsOligo 2
Activated dsOligo 1
Cys
Cys
dsOligo-protein-dsOligo
Chimera
dsOligo 2
dsOligo 1

**Fig. S2**

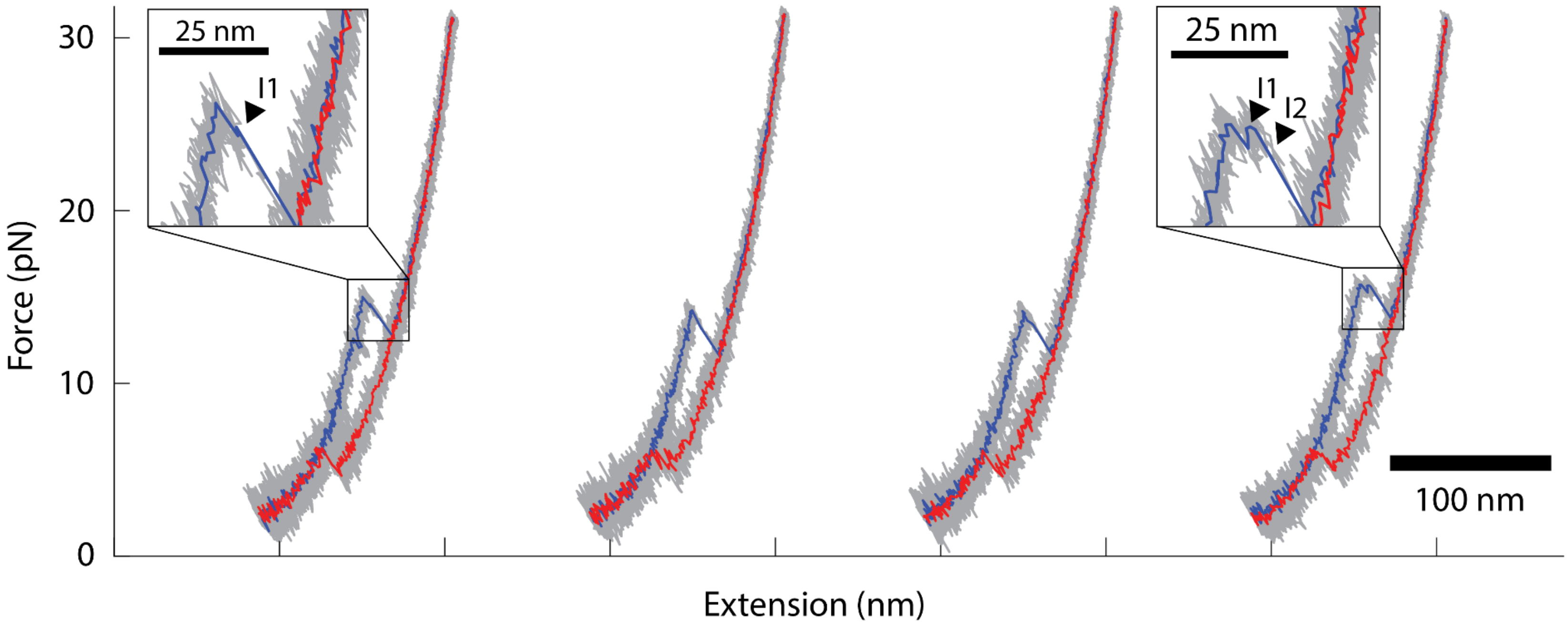


**Fig. S3**

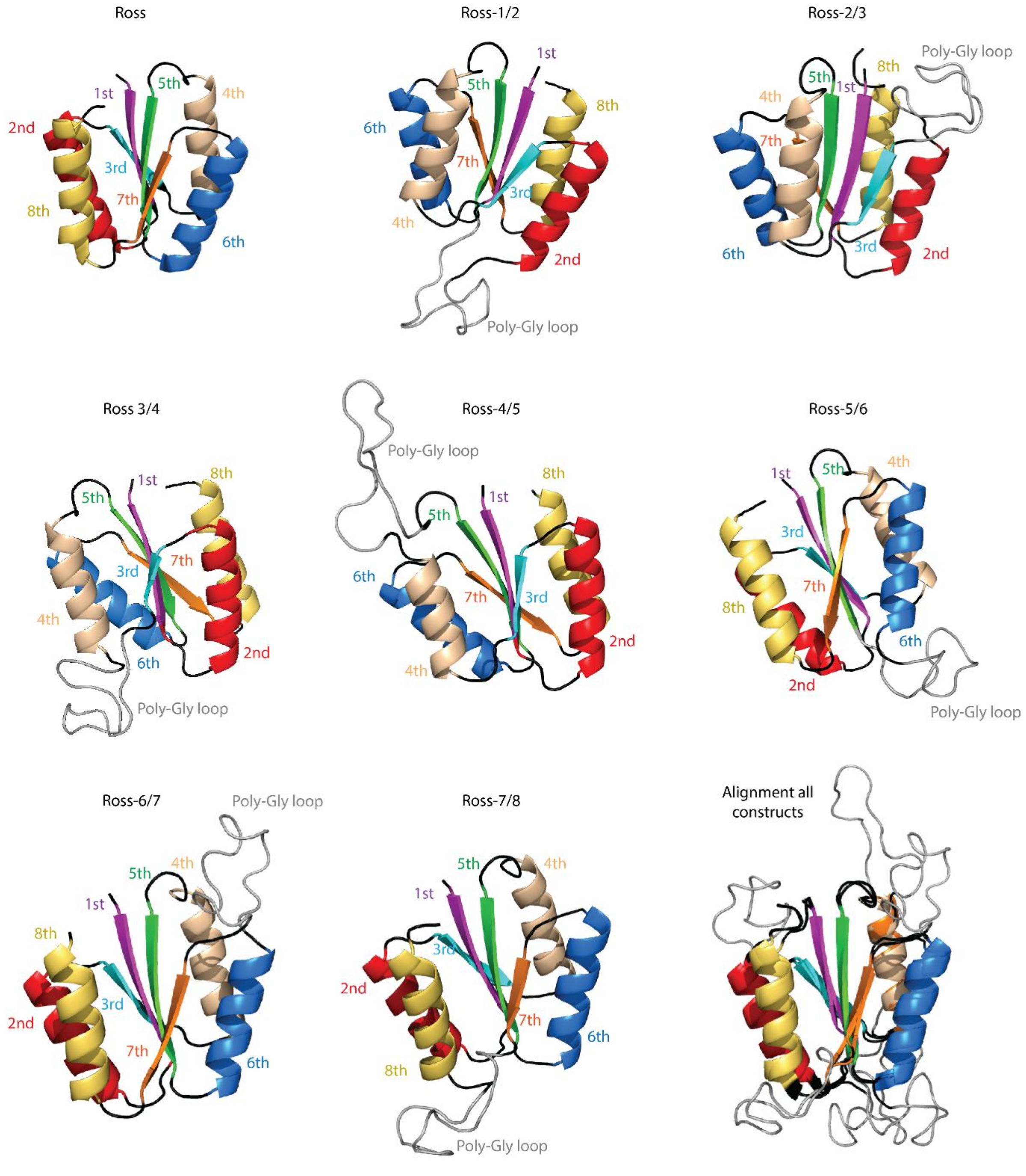

Ross
1st
5th
4th
2nd
3rd
7th
8th
6th
Ross-1/2
1st
5th
6th
8th
7th
3rd
4th
2nd
Poly-Gly loop
Ross-2/3
Poly-Gly loop
5th
8th
1st
4th
7th
6th
3rd
2nd
Ross 3/4
8th
1st
5th
7th
3rd
4th
2nd
6th
Poly-Gly loop
Ross-4/5
Poly-Gly loop
8th
1st
5th
6th
7th
3rd
2nd
4th
Ross-5/6
4th
5th
1st
3rd
7th
8th
6th
2nd
Poly-Gly loop
Ross-6/7
Poly-Gly loop
5th
4th
1st
8th
3rd
2nd
7th
6th
Ross-7/8
5th
4th
1st
3rd
2nd
7th
6th
8th
Poly-Gly loop
Alignment all
constructs


**Fig. S4**

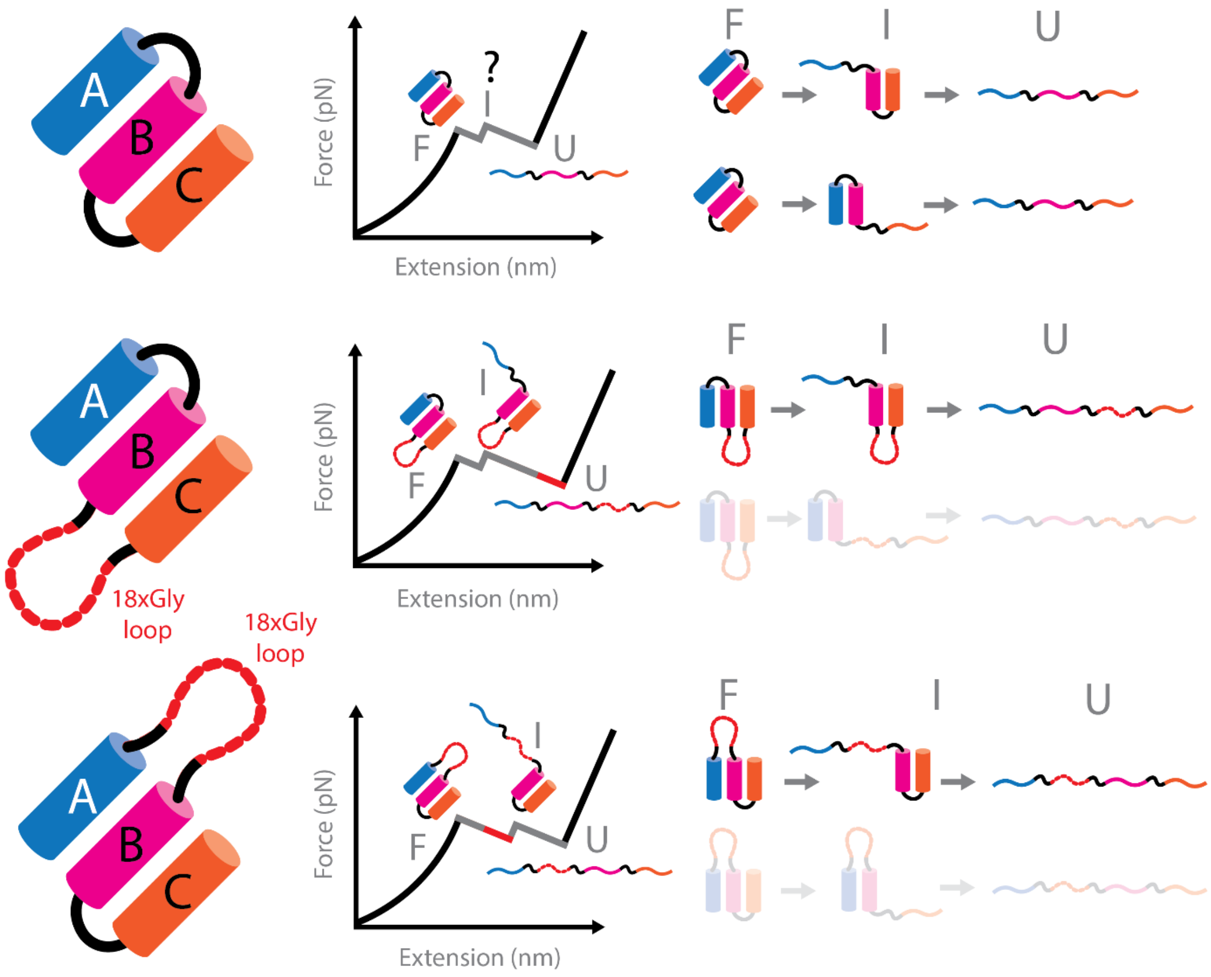
A
B
C
?
I
F
U
Force (pN)
Extension (nm)
F
I
U
A
B
C
18xGly
loop
I
F
U
Force (pN)
Extension (nm)
F
I
U
18xGly
loop
A
B
C
I
F
U
Force (pN)
Extension (nm)
F
I
U

**Fig. S5**

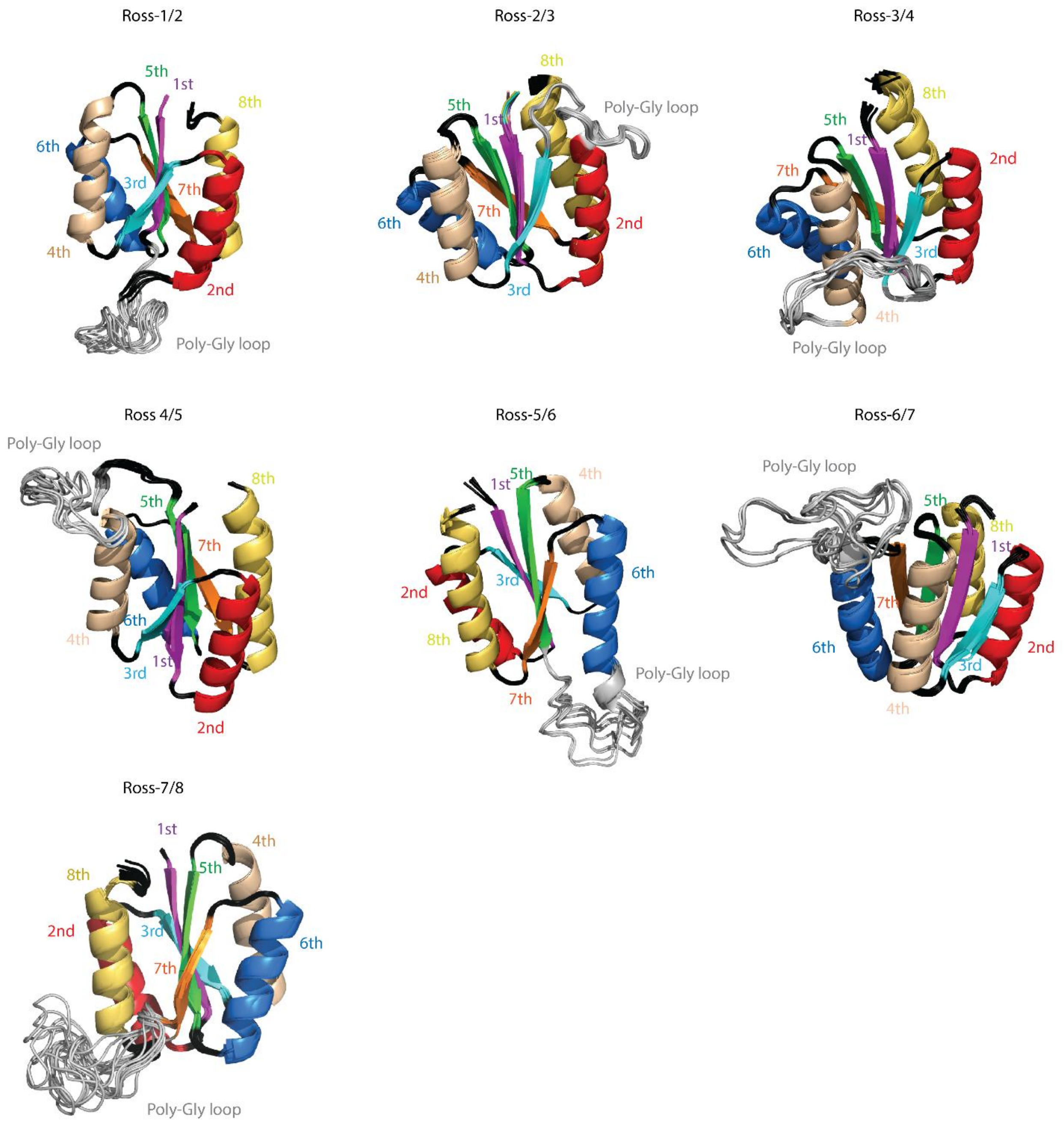
Ross-1/2
5th
1st
8th
6th
3rd
7th
4th
2nd
Poly-Gly loop
Ross-2/3
8th
Poly-Gly loop
5th
1st
6th
7th
2nd
4th
3rd
Ross-3/4
8th
5th
1st
2nd
7th
6th
3rd
4th
Poly-Gly loop
Ross 4/5
Poly-Gly loop
8th
5th
7th
6th
4th
1st
3rd
2nd
Ross-5/6
5th
4th
1st
6th
3rd
2nd
8th
Poly-Gly loop
7th
Ross-6/7
Poly-Gly loop
5th
8th
1st
7th
6th
2nd
3rd
4th
Ross-7/8
1st
4th
5th
8th
2nd
3rd
6th
7th
Poly-Gly loop

**Fig. S6**

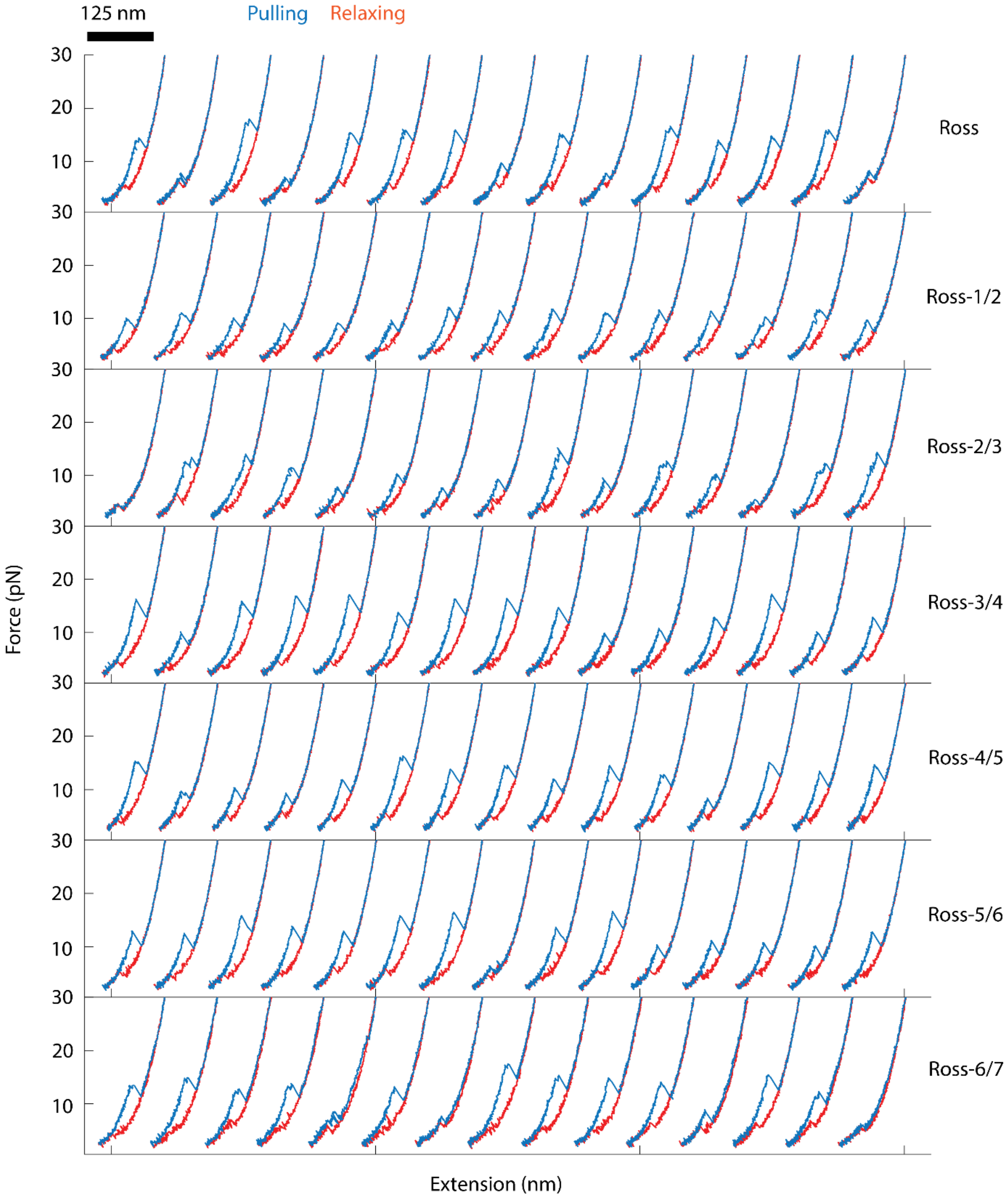
125 nm
Pulling
Relaxing
30
20
10
Ross
Ross-1/2
Ross-2/3
Ross-3/4
Ross-4/5
Ross-5/6
Ross-6/7
Force (pN)
Extension (nm)

**Fig. S7**

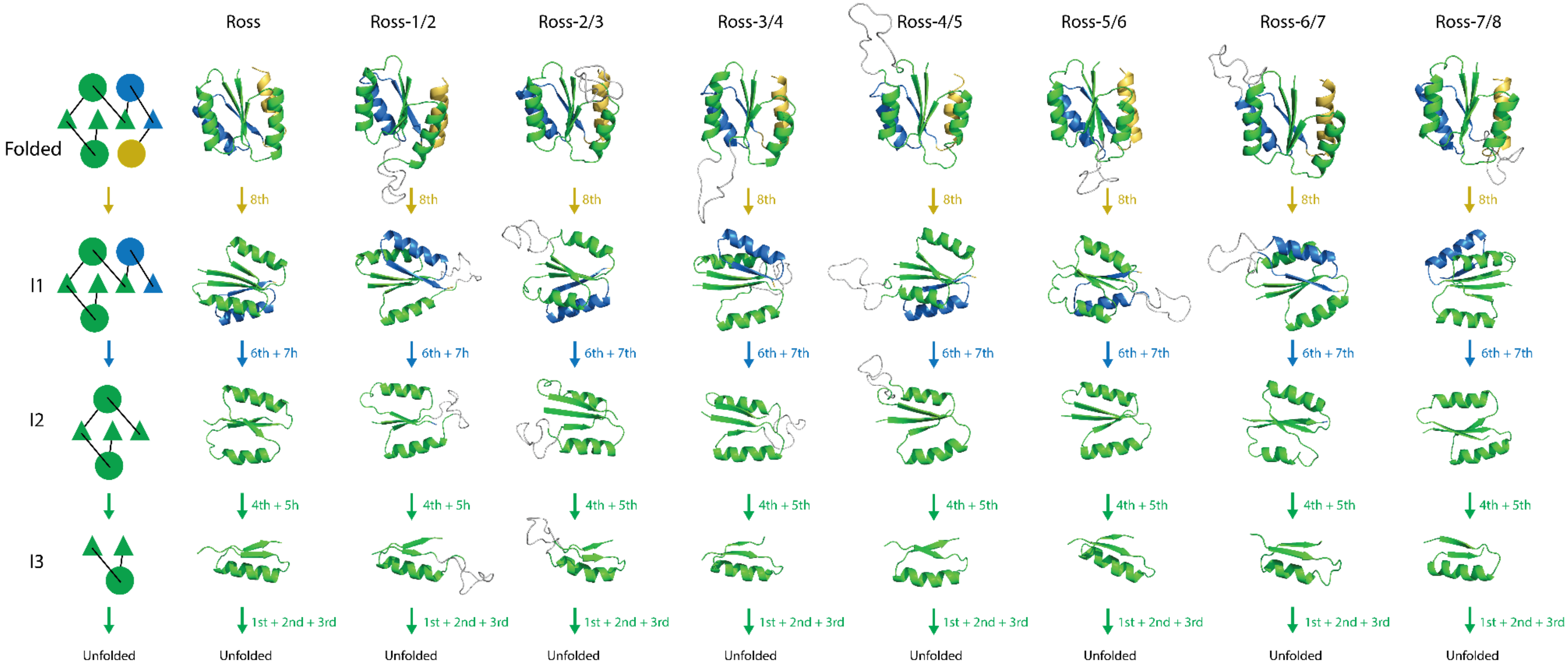

Ross
Ross-1/2
Ross-2/3
Ross-3/4
Ross-4/5
Ross-5/6
Ross-6/7
Ross-7/8
Folded
8th
I1
6th + 7h
6th + 7th
I2
4th + 5h
4th + 5th
I3
1st + 2nd + 3rd
Unfolded


**Fig. S8**

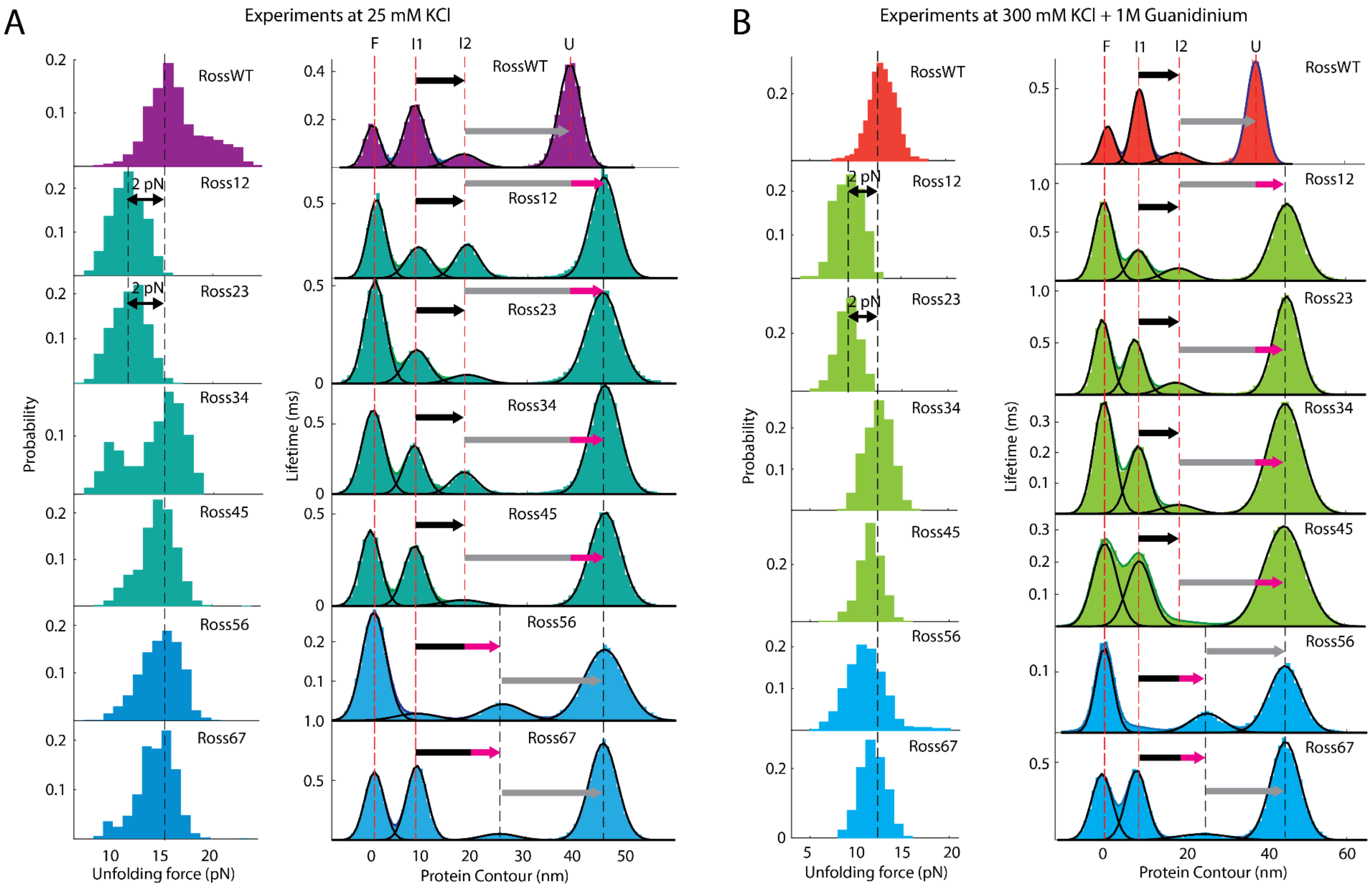
A
Experiments at 25 mM KCl
B
Experiments at 300 mM KCl + 1M Guanidinium
F
I1
I2
U
RossWT
Ross12
Ross23
Ross34
Ross45
Ross56
Ross67
2 pN
Probability
Lifetime (ms)
Unfolding force (pN)
Protein Contour (nm)

**Fig. S9**

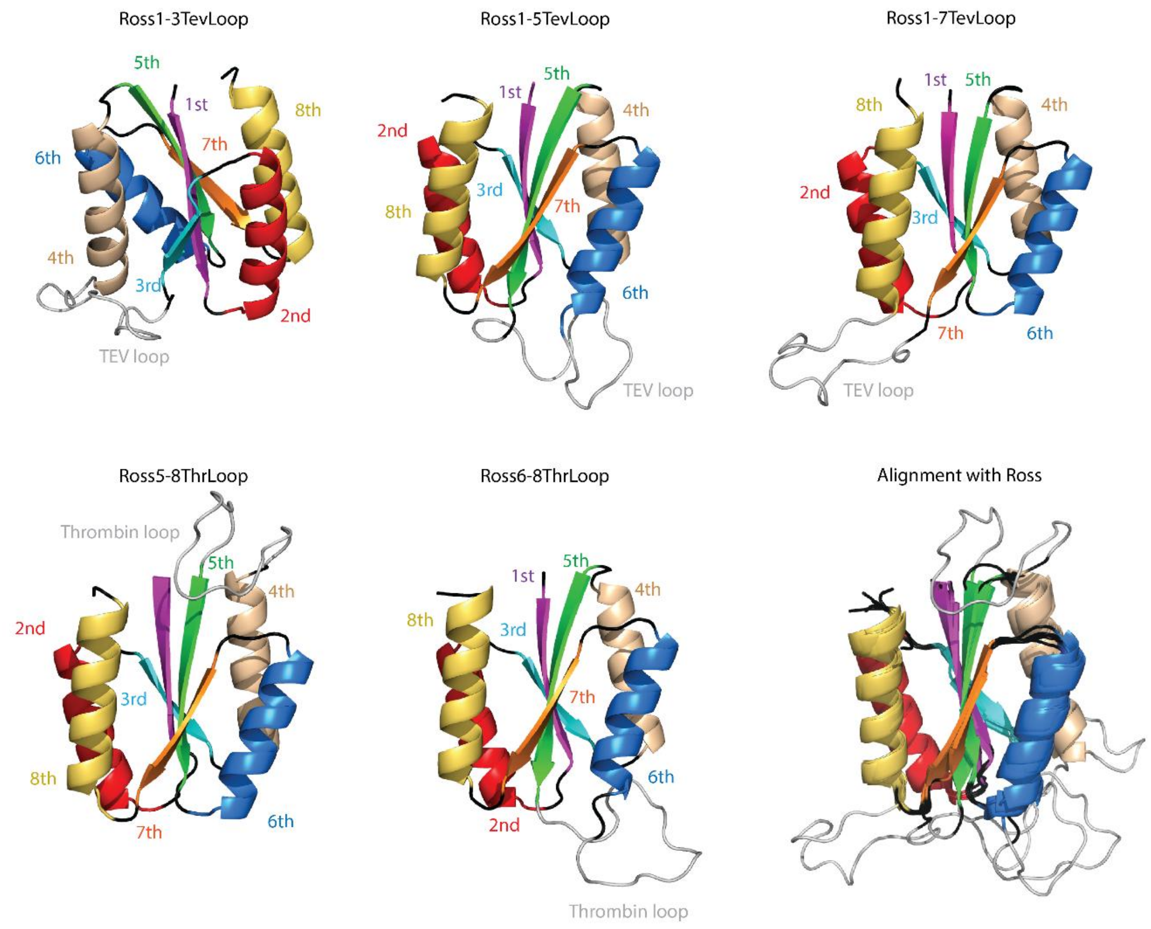
Ross1-3TevLoop
5th
1st
8th
7th
6th
4th
3rd
2nd
TEV loop
Ross1-5TevLoop
1st
5th
4th
2nd
3rd
8th
7th
6th
TEV loop
Ross1-7TevLoop
1st
5th
8th
4th
2nd
3rd
7th
6th
TEV loop
Ross5-8ThrLoop
Thrombin loop
5th
4th
2nd
3rd
8th
7th
6th
Ross6-8ThrLoop
1st
5th
4th
8th
3rd
7th
6th
2nd
Thrombin loop
Alignment with Ross

**Fig. S10**

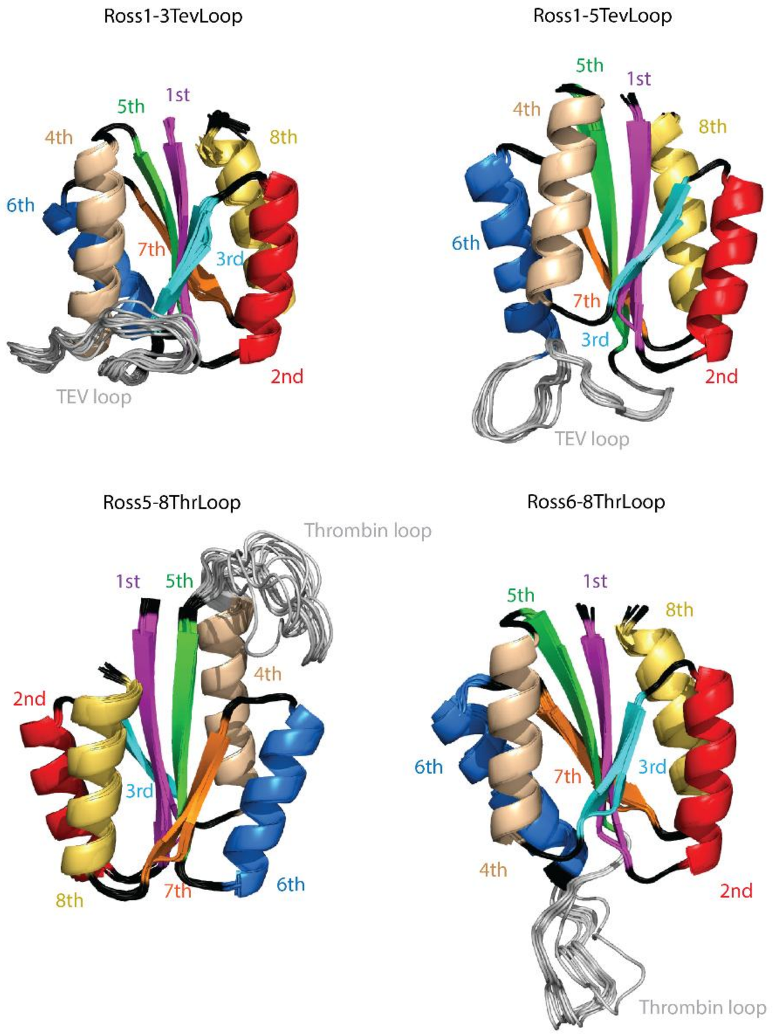

Ross1-3TevLoop
5th
1st
4th
8th
6th
7th
3rd
2nd
TEV loop
Ross1-5TevLoop
5th
1st
4th
8th
6th
7th
3rd
2nd
TEV loop
Ross1-7TevLoop
1st
5th
4th
2nd
3rd
6th
8th
7th
TEV loop
Ross5-8ThrLoop
Thrombin loop
1st
5th
4th
2nd
3rd
8th
7th
6th
Ross6-8ThrLoop
5th
1st
8th
6th
7th
3rd
4th
2nd
Thrombin loop


**Fig. S11**

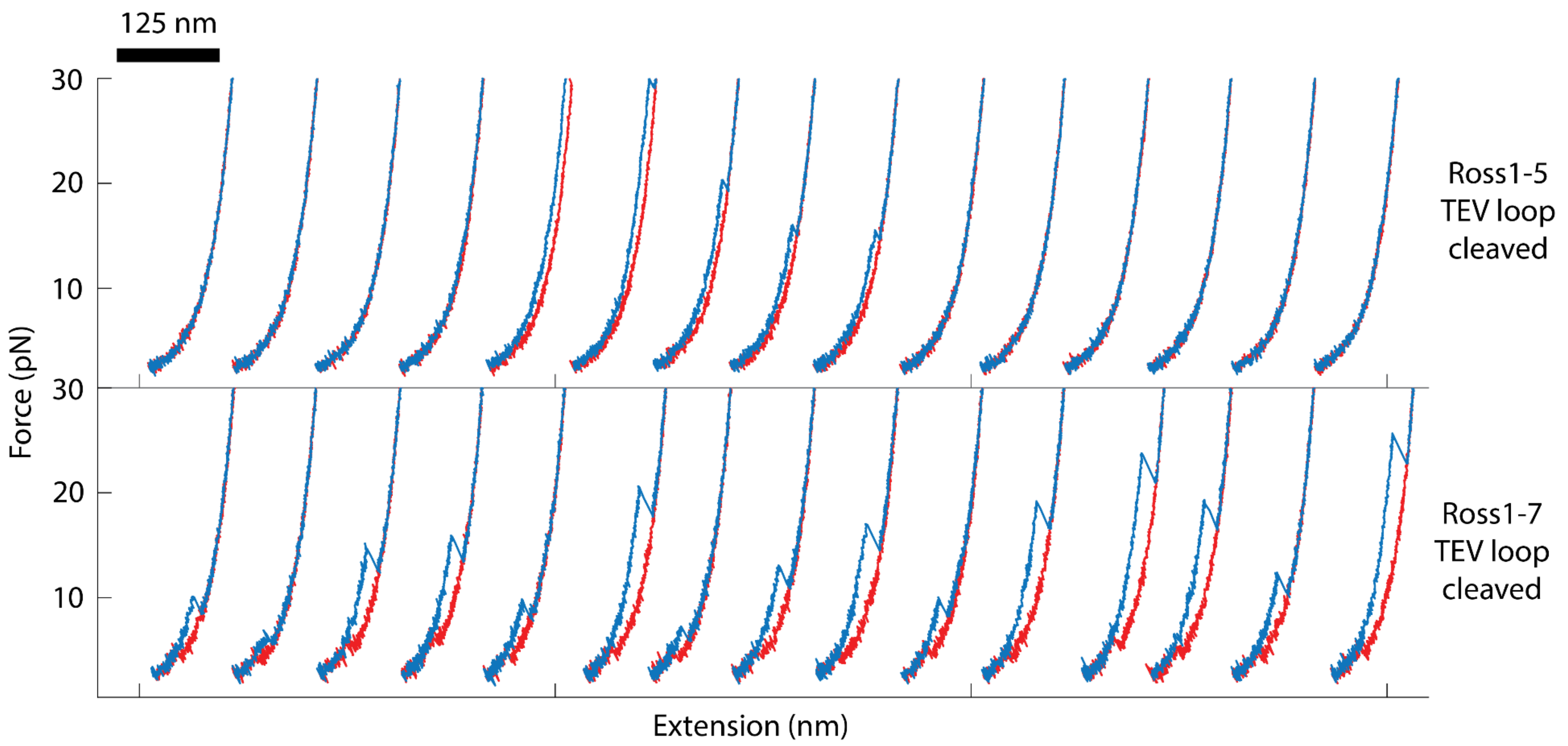
125 nm
30
20
10
30
20
10
Force (pN)
Extension (nm)
Ross1-5 TEV loop cleaved
Ross1-7 TEV loop cleaved

**Fig. S12**

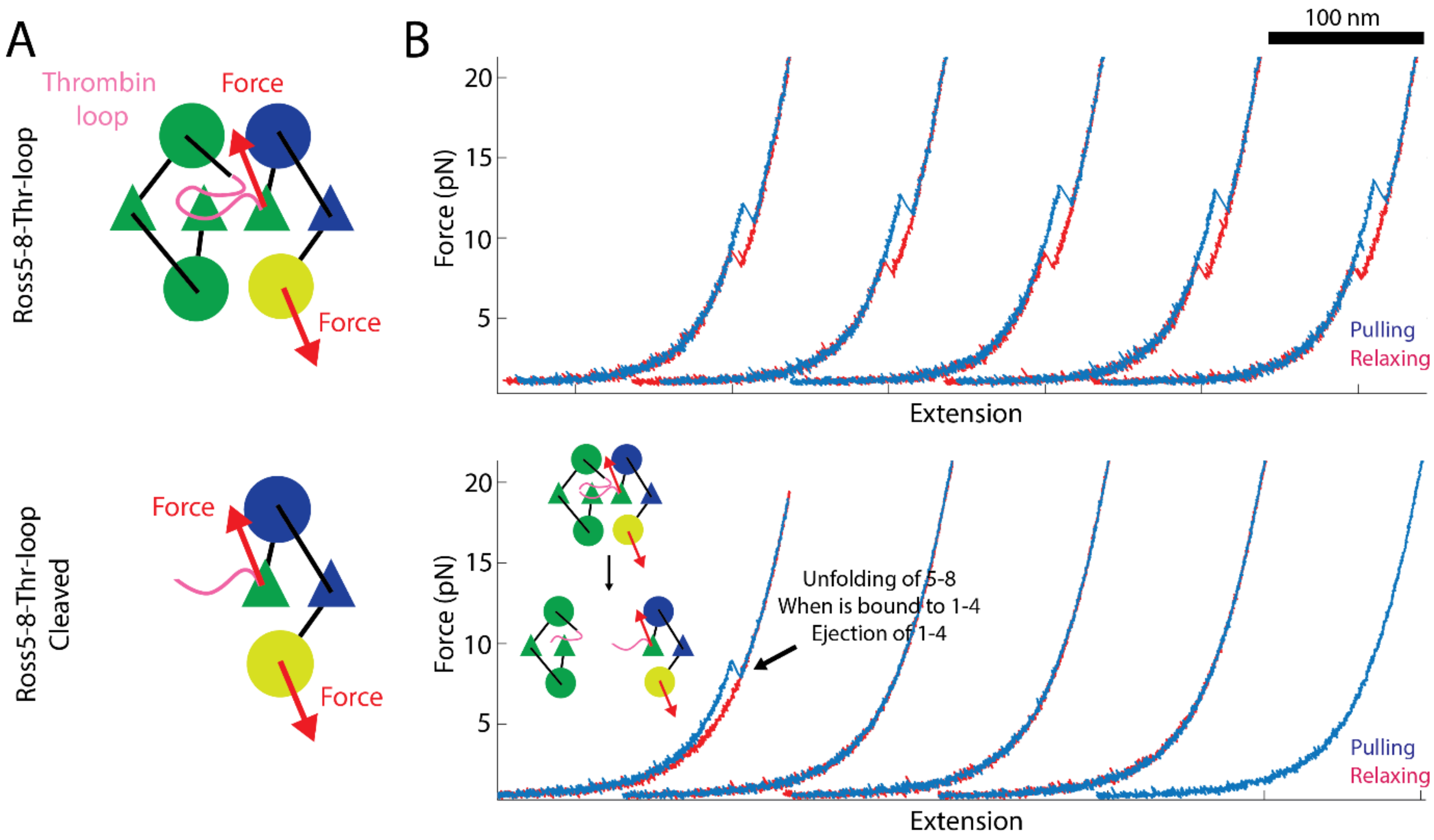
A
Thrombin loop
Force
Force
Ross5-8-Thr-loop
Ross5-8-Thr-loop Cleaved
Force
Force
B
100 nm
Force (pN)
20
15
10
5
Extension
Pulling
Relaxing
Unfolding of 5-8
When is bound to 1-4
Ejection of 1-4

**Fig. S13**

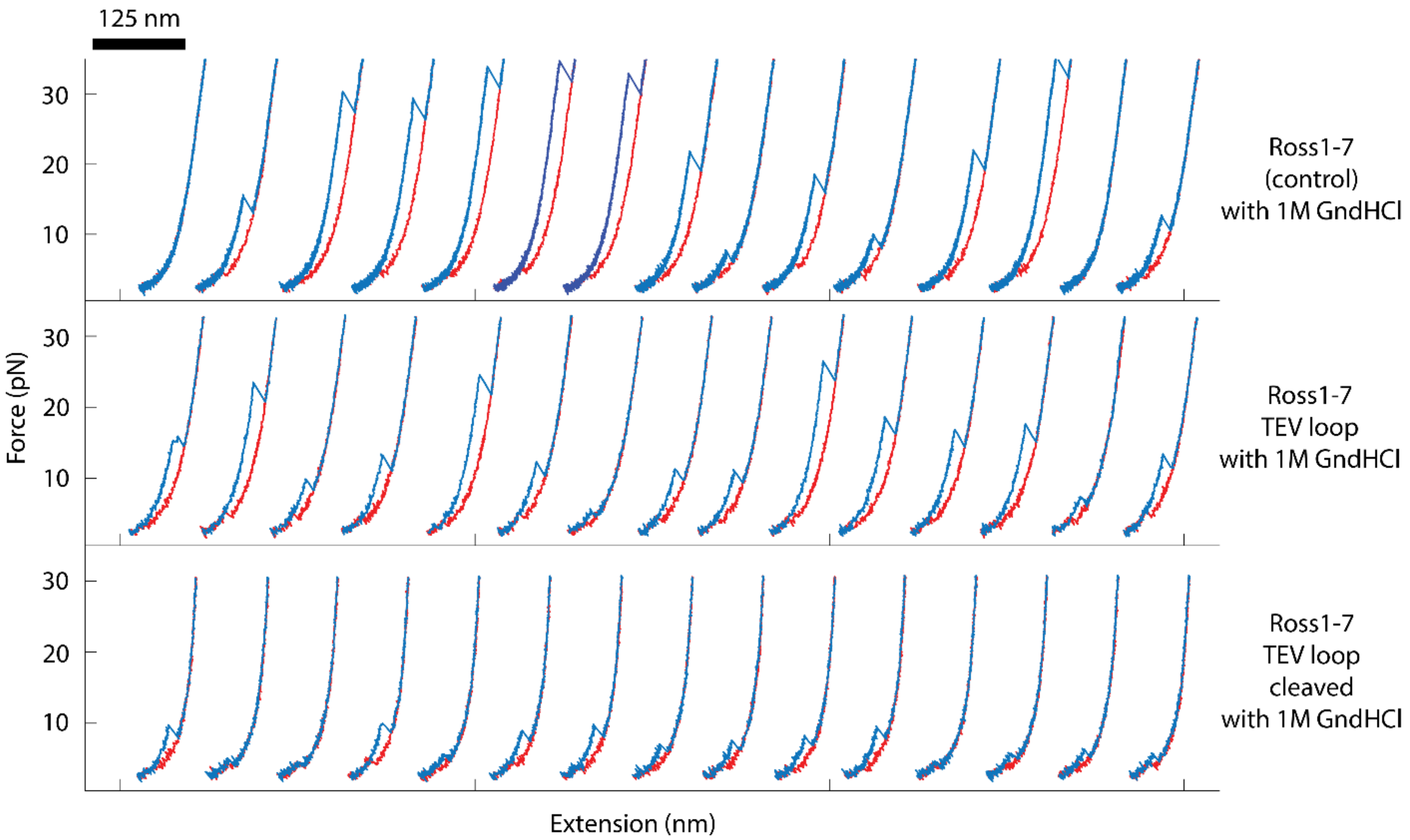
125 nm
30
20
10
Force (pN)
Extension (nm)
Ross1-7
(control)
with 1M GndHCl
Ross1-7
TEV loop
with 1M GndHCl
Ross1-7
TEV loop
cleaved
with 1M GndHCl

**Fig. S14**

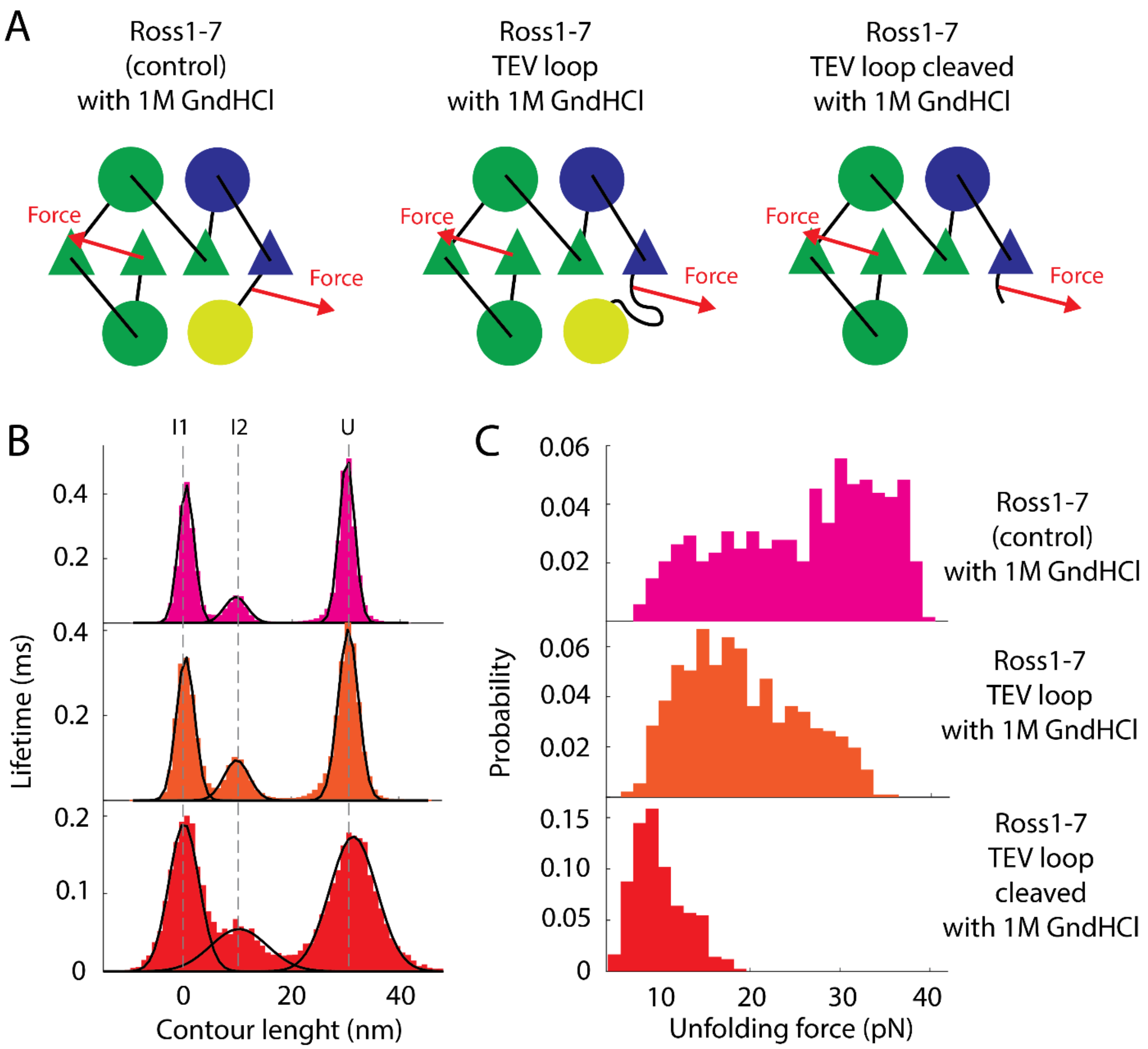
A
Ross1-7
(control)
with 1M GndHCl
Ross1-7
TEV loop
with 1M GndHCl
Ross1-7
TEV loop cleaved
with 1M GndHCl
Force
Force
Force
Force
Force
Force
B
I1
I2
U
0.4
0.2
0.4
0.2
0.2
0.1
0
Lifetime (ms)
0
20
40
Contour lenght (nm)
C
0.06
0.04
0.02
0.06
0.04
0.02
0.15
0.10
0.05
0
Probability
10
20
30
40
Unfolding force (pN)
Ross1-7
(control)
with 1M GndHCl
Ross1-7
TEV loop
with 1M GndHCl
Ross1-7
TEV loop
cleaved
with 1M GndHCl

**Fig. S15**

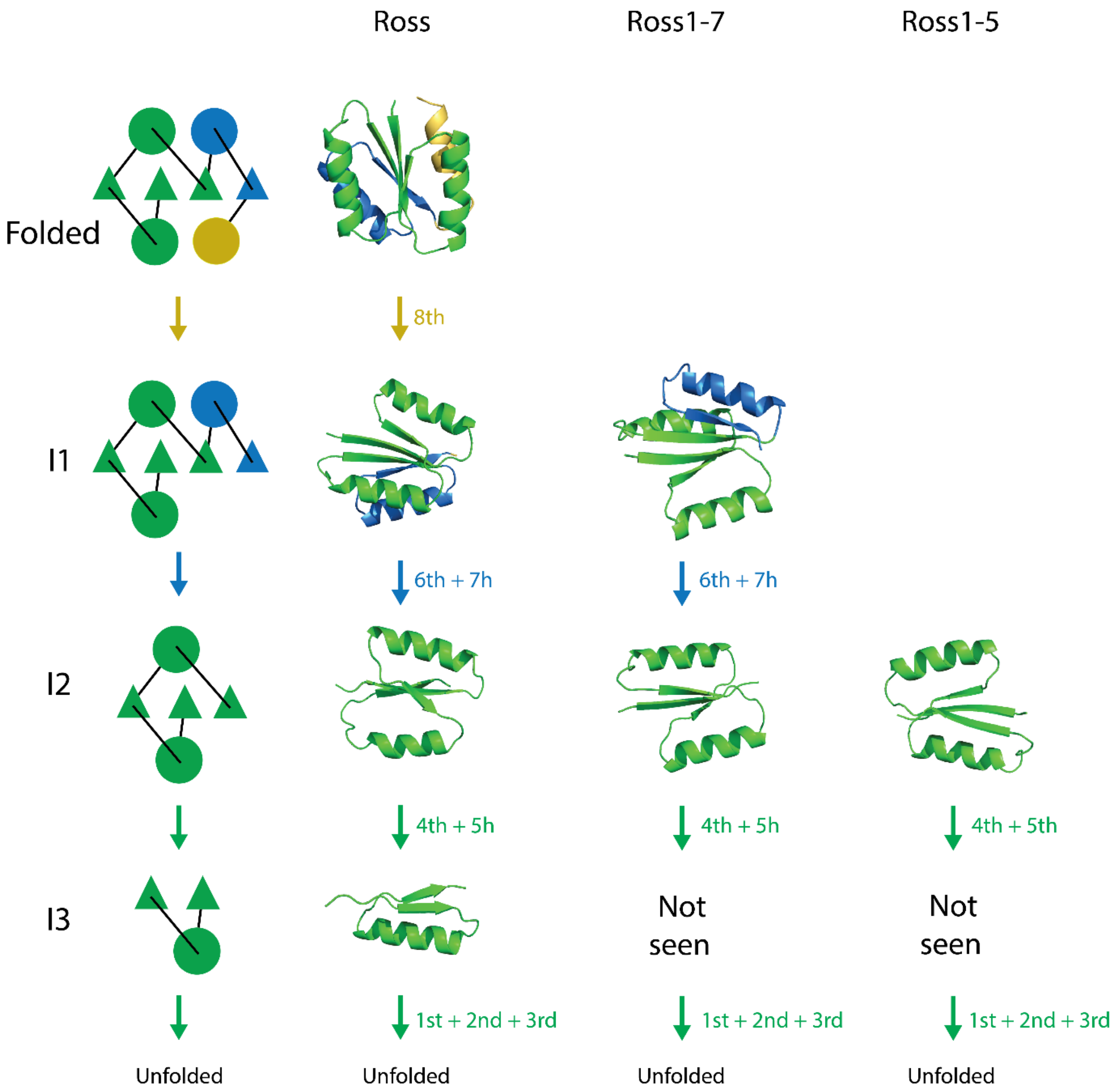
Ross
Ross1-7
Ross1-5
Folded
8th
I1
6th + 7h
6th + 7h
I2
4th + 5h
4th + 5h
4th + 5th
I3
Not seen
Not seen
1st + 2nd + 3rd
1st + 2nd + 3rd
1st + 2nd + 3rd
Unfolded
Unfolded
Unfolded
Unfolded

**Fig. S16**

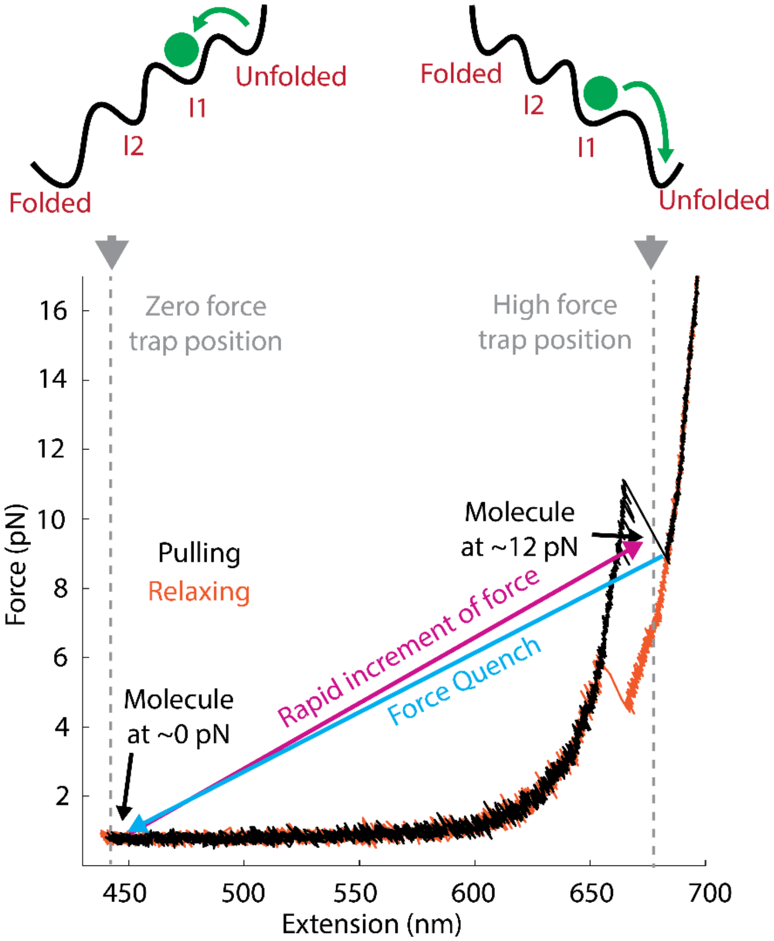
Folded
I2
I1
Unfolded
Folded
I2
I1
Unfolded
Zero force
trap position
High force
trap position
Molecule
at ~12 pN
Pulling
Relaxing
Rapid increment of force
Force Quench
Molecule
at ~0 pN
Force (pN)
16
14
12
10
8
6
4
2
450
500
550
600
650
700
Extension (nm)

**Fig. S17**

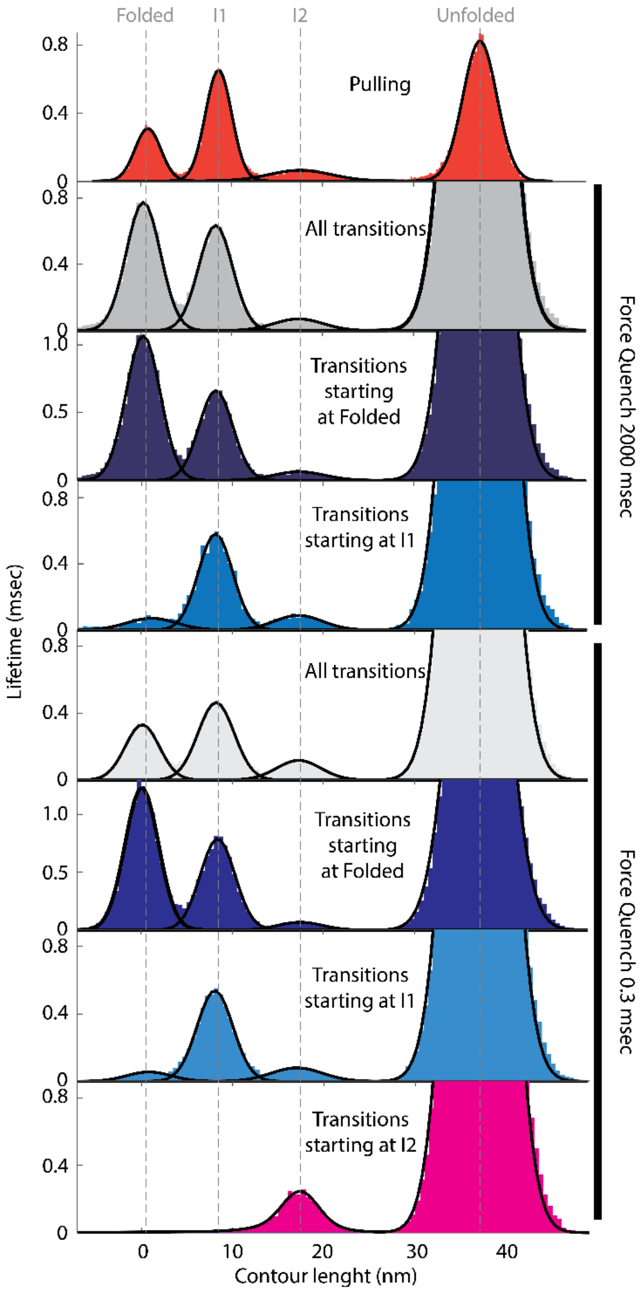

Folded
I1
I2
Unfolded
Pulling
All transitions
Transitions starting at Folded
Transitions starting at I1
All transitions
Transitions starting at Folded
Transitions starting at I1
Transitions starting at I2
Force Quench 2000 msec
Force Quench 0.3 msec
Lifetime (msec)
Contour lenght (nm)
0
0.4
0.8
0.5
1.0
10
20
30
40


**Fig. S18**

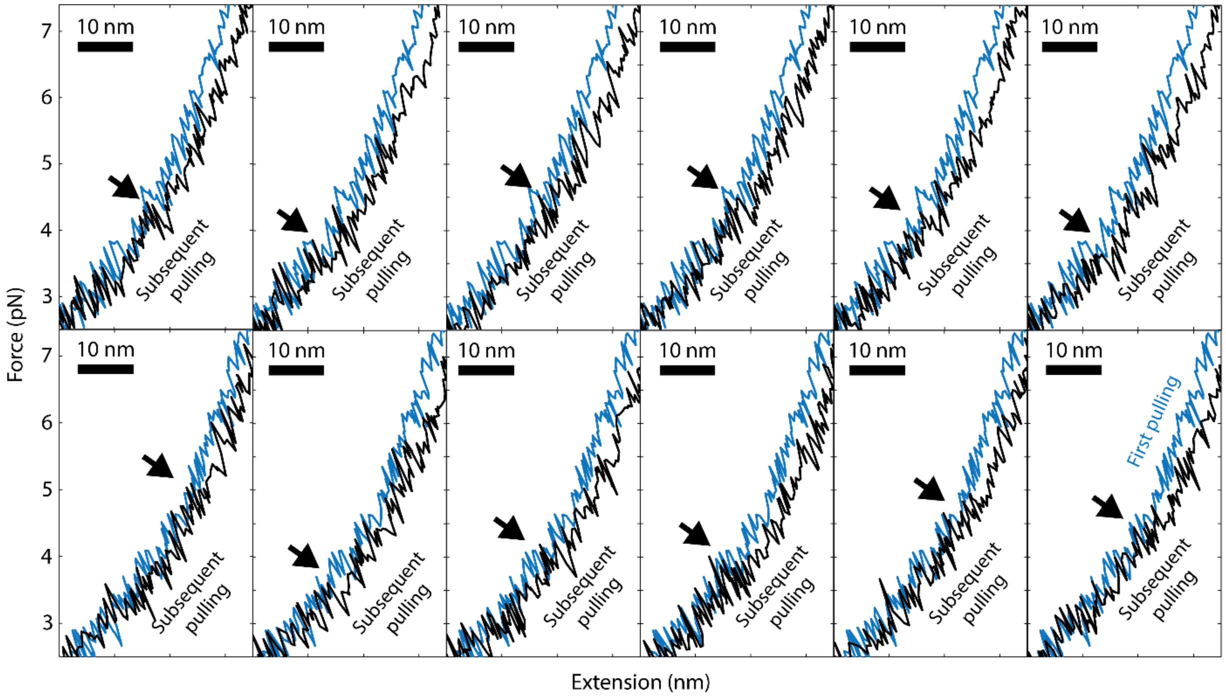
Force (pN)
Extension (nm)
10 nm
Subsequent pulling
First pulling
3
4
5
6
7

**Fig. S19**

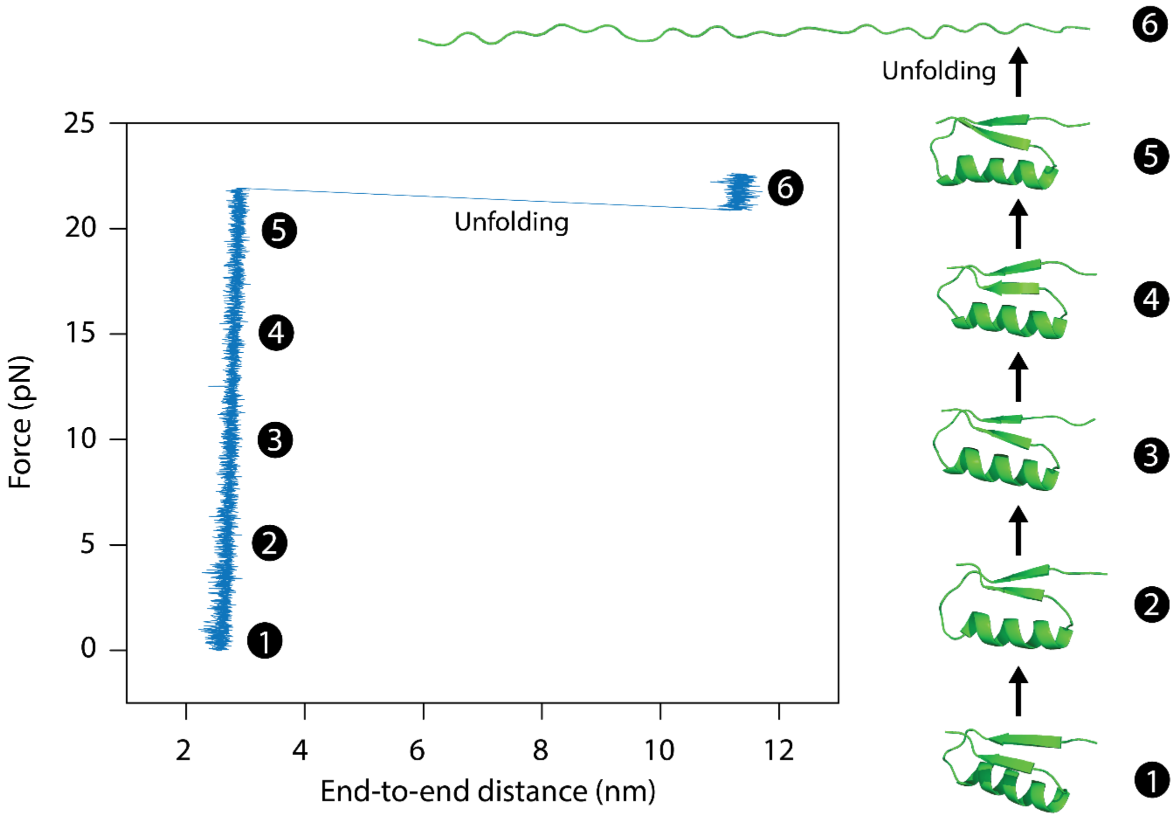
25
20
15
10
5
0
Force (pN)
2
4
6
8
10
12
End-to-end distance (nm)
Unfolding
Unfolding
1
2
3
4
5
6

**Fig. S20**

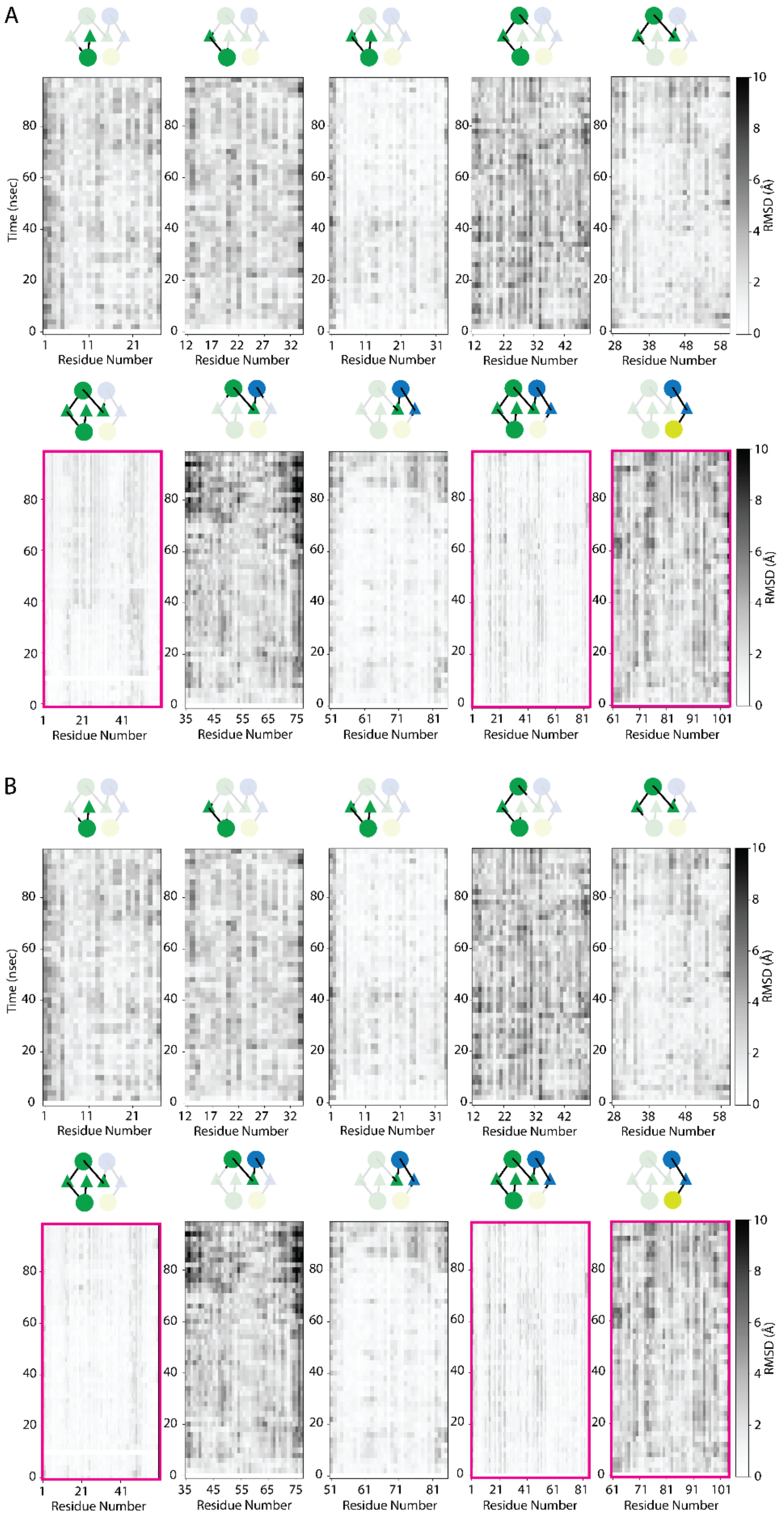


**Fig. S21**

**Table S1:** List of protein variants of Ross

| Protein | Sequence |
|---|---|
| Ross | MACKMLLYVLIISNDKKLIEEARKMAEKANLELRTVKTEDELKKYLEE<br>FRKESQNIKVLILVSNDEELDKAKELAQKMEIDVRTRKVTSPDEAKRW<br>IKEFSEEGGSKCLEHHHHHH |
| Ross-1/2 | MACKMLLYVLIIGGGGGGGGGGGGGGGGGGSNDKKLIEEARKMAE<br>KANLELRTVKTEDELKKYLEEFRKESQNIKVLILVSNDEELDKAKELA<br>QKMEIDVRTRKVTSPDEAKRWIKEFSEEGGSKCLEHHHHHH |
| Ross-2/3 | MACKMLLYVLIISNDKKLIEEARKMAEKANGGGGGGGGGGGGGGGG<br>GGLELRTVKTEDELKKYLEEFRKESQNIKVLILVSNDEELDKAKELAQ<br>KMEIDVRTRKVTSPDEAKRWIKEFSEEGGSKCLEHHHHHH |
| Ross-3/4 | MACKMLLYVLIISNDKKLIEEARKMAEKANLELRTVGGGGGGGGGG<br>GGGGGGGGKTEDELKKYLEEFRKESQNIKVLILVSNDEELDKAKELA<br>QKMEIDVRTRKVTSPDEAKRWIKEFSEEGGSKCLEHHHHHH |
| Ross-4/5 | MACKMLLYVLIISNDKKLIEEARKMAEKANLELRTVKTEDELKKYLEE<br>FRKGGGGGGGGGGGGGGGGGGESQNIKVLILVSNDEELDKAKELAQ<br>KMEIDVRTRKVTSPDEAKRWIKEFSEEGGSKCLEHHHHHH |
| Ross-5/6 | MACKMLLYVLIISNDKKLIEEARKMAEKANLELRTVKTEDELKKYLEE<br>FRKESQNIKVLILVGGGGGGGGGGGGGGGGGGSNDEELDKAKELAQ<br>KMEIDVRTRKVTSPDEAKRWIKEFSEEGGSKCLEHHHHHH |
| Ross-6/7 | MACKMLLYVLIISNDKKLIEEARKMAEKANLELRTVKTEDELKKYLEE<br>FRKESQNIKVLILVSNDEELDKAKELAQKMGGGGGGGGGGGGGGGG<br>GGEIDVRTRKVTSPDEAKRWIKEFSEEGGSKCLEHHHHHH |
| Ross-7/8 | MACKMLLYVLIISNDKKLIEEARKMAEKANLELRTVKTEDELKKYLEE<br>FRKESQNIKVLILVSNDEELDKAKELAQKMEIDVRTRKGGGGGGGGG<br>GGGGGGGGGVTSPDEAKRWIKEFSEEGGSKCLEHHHHHH |
| Ross1-7-TevLoop | MACKMLLYVLIISNDKKLIEEARKMAEKANLELRTVKTEDELKKYLEE<br>FRKESQNIKVLILVSNDEELDKAKELAQKMEIDVRTRKVTCGGGGGEN<br>LYFQGGGGGGGSPDEAKRWIKEFSEEGGSKLEHHHHHH |
| Ross1-3 TevLoop | MACKMLLYVLIISNDKKLIEEARKMAEKANLELRTVKCGGGGGENLY<br>FQ//GGGGGGGTEDELKKYLEEFRKESQNIKVLILVSNDEELDKAKELA<br>QKMEIDVRTRKVTSPDEAKRWIKEFSEEGGSKSLEHHHHHH |
| Ross1-5-TevLoop | MACKMLLYVLIISNDKKLIEEARKMAEKANLELRTVKTEDELKKYLEE<br>FRKESQNIKVLILVSCGGGGGENLYFQ//GGGGGGGNDEELDKAKELA<br>QKMEIDVRTRKVTSPDEAKRWIKEFSEEGGSKLEHHHHHH |
| Ross6-8-ThrLoop | MASKMLLYVLIISNDKKLIEEARKMAEKANLELRTVKTEDELKKYLEE<br>FRKESQNIKVLILVGGTEGSLVPRGS//GESGGCNDEELDKAKELAQKM<br>EIDVRTRKVTSPDEAKRWIKEFSEEGGSKCLEHHHHHH |
| Ross5-8-ThrLoop | MASKMLLYVLIISNDKKLIEEARKMAEKANLELRTVKTEDELKKYLEE<br>FRKESQGGTEGSLVPRGS//GESGGCNIKVLILVSNDEELDKAKELAQK<br>MEIDVRTRKVTSPDEAKRWIKEFSEEGGSKCLEHHHHHH |

*Each secondary elements are bold color-coded, loops are in black bold, cys are in black bold, Gly loops are in grey bold. Other elements such as HisTag are in black.

**Table S2:** Contour length values of all folded, unfolded and intermediates states during PM experiments of Ross

| Experiment | Path | Folded | I1 | I2 | Unfolded | n |
|---|---|---|---|---|---|---|
| **Pull/relax** | Unfolding | 0.81 ± 0.06 | 8.64 ± 0.06 | 17.67 ± 0.04 | 37.59 ± 0.03 | 823 |
| **PM mode** | Folding | 0.38 ± 0.04 | 8.13 ± 0.03 | 17.83 ± 0.05 | 37.47 ± 0.25 | 1156 |
| | Unfolding | 0.41 ± 0.04 | 8.46 ± 0.04 | 17.24 ± 0.05 | 37.41 ± 0.29 | 1159 |
| | Incomplete folding | --- | 8.86 ± 1.50 | 17.79 ± 0.04 | 36.08 ± 0.16 | 2262 |

**Table S3:** Contour length values of all folded, unfolded and intermediates states during pulling/relaxing experiments of all Ross constructs and its variants

| Cond. | Construct | Folded | I1 | I2 | Unfolded | n |
|---|---|---|---|---|---|---|
| **300 mM KCl** | **Ross** | 0.81 ± 0.06 | 8.64 ± 0.06 | 17.67 ± 0.04 | 37.59 ± 0.03 | 823 |
| | **Ross-1/2** | 0.91 ± 0.04 | 8.38 ± 0.05 | 17.85 ± 0.09 | 42.84 ± 0.12 | 527 |
| | **Ross-2/3** | 1.30 ± 0.10 | 7.97 ± 0.10 | 17.60 ± 0.16 | 43.49 ± 0.13 | 439 |
| | **Ross-3/4** | 0.50 ± 0.05 | 8.43 ± 0.06 | 17.98 ± 0.07 | 42.24 ± 0.11 | 756 |
| | **Ross-4/5** | 0.98 ± 0.06 | 9.04 ± 0.06 | 17.37 ± 0.04 | 43.69 ± 0.05 | 930 |
| | **Ross-5/6** | 0.58 ± 0.02 | 8.26 ± 0.03 | 24.43 ± 0.05 | 43.33 ± 1.01 | 850 |
| | **Ross-6/7** | 0.91 ± 0.12 | 8.94 ± 0.12 | 24.42 ± 0.04 | 43.71 ± 0.04 | 970 |
| | **Ross-1-7** | ND | ND | ND | ND | --- |
| | **Ross-1-7 TEVloop** | ND | ND | ND | ND | --- |
| | **Ross-1-7 TEVloop*** | --- | 7.27 + 0.04 | 16.95 ± 0.04 | 38.38 ± 0.05 | 636 |
| | **Ross-1-5** | ND | ND | ND | ND | --- |
| | **Ross-1-5 TEVloop** | ND | ND | ND | ND | --- |
| | **Ross-1-5 TEVloop*** | --- | --- | 16.44 ± 0.13 | 38.38 ± 0.13 | 18 |
| | **Ross-6-8 Thrombin*** | 1.22 ± 0.07 | 9.36 ± 0.10 | 16.58 ± 0.23 | --- | 213 |
| **25 mM KCl** | **Ross** | -0.11 ± 0.04 | 8.10 ± 0.04 | 17.23 ± 0.05 | 37.97 ± 0.07 | 746 |
| | **Ross-1/2** | 0.7 ± 0.07 | 8.82 ± 0.08 | 18.28 ± 0.09 | 44.76 ± 0.22 | 423 |
| | **Ross-2/3** | 0.05 ± 0.07 | 8.06 ± 0.08 | 17.74 ± 0.10 | 45.13 ± 0.12 | 469 |
| | **Ross-3/4** | 0.42 ± 0.06 | 8.52 ± 0.06 | 18.32 ± 0.07 | 44.73 ± 0.21 | 703 |
| | **Ross-4/5** | -0.46 ± 0.05 | 8.14 ± 0.05 | 17.52 ± 0.04 | 45.18 ± 0.07 | 731 |
| | **Ross-5/6** | 0.22 ± 0.05 | 8.57 ± 0.05 | 25.31 ± 0.05 | 45.07 ± 1.29 | 734 |
| | **Ross-6/7** | 0.36 ± 0.04 | 8.62 ± 0.04 | 25.74 ± 0.05 | 45.00 ± 0.04 | 690 |
| **300 mM KCl, 1M GndHCl** | **Ross** | 1.07 ± 0.08 | 8.87 ± 0.08 | 18.23 ± 0.04 | 37.92 ± 0.05 | 835 |
| | **Ross-1/2** | 0.10 ± 0.08 | 8.55 ± 0.08 | 18.50 ± 0.14 | 45.42 ± 0.23 | 215 |
| | **Ross-2/3** | -0.30 ± 0.08 | 7.72 ± 0.09 | 17.83 ± 0.14 | 45.36 ± 0.13 | 411 |
| | **Ross-3/4** | 0.11 ± 0.06 | 8.61 ± 0.06 | 18.60 ± 0.05 | 44.97 ± 0.11 | 459 |
| | **Ross-4/5** | 0.07 ± 0.12 | 8.62 ± 0.14 | 18.31 ± 0.11 | 44.63 ± 0.15 | 607 |
| | **Ross-5/6** | 0.17 ± 0.08 | 8.97 ± 0.14 | 25.54 ± 0.11 | 44.73 ± 2.33 | 452 |
| | **Ross-6/7** | -0.46 ± 0.09 | 8.09 ± 0.10 | 24.53 ± 0.09 | 44.84 ± 0.09 | 454 |
| | **Ross-1-7** | --- | 8.37 ± 0.05 | 17.56 ± 0.07 | 37.13 ± 0.07 | 488 |
| | **Ross-1-7 TEVloop** | --- | 8.30 ± 0.06 | 17.82 ± 0.09 | 37.62 ± 0.07 | 651 |
| | **Ross-1-7 TEVloop*** | --- | 7.95 ± 0.17 | 18.15 ± 0.16 | 38.32 ± 0.12 | 301 |
| | **Ross-1-5[+]** | ND | ND | ND | ND | --- |
| | **Ross-1-5 TEVloop[+]** | ND | ND | ND | ND | --- |
| | **Ross-1-5 TEVloop*[+]** | ND | ND | ND | ND | --- |

* Cleaved with the respective protease.
ND = not determined

**Table S4:** Contour length values of all folded, unfolded and intermediates states from RFQ experiments

| Condition | Transitions | Folded | I1 | I2 | Unfolded | n |
|---|---|---|---|---|---|---|
| **2000 msec** | All | 0.26 ± 0.18 | 8.22 ± 0.18 | 17.34 ± 0.37 | 37.12 ± 0.34 | 382 |
| | Starting at folded | 0.24 ± 0.21 | 8.25 ± 0.21 | 17.40 ± 0.47 | 37.11 ± 0.41 | 270 |
| | Starting at I1 | --- | 8.15 ± 0.33 | 17.28 ± 0.65 | 37.14 ± 0.64 | 110 |
| | Starting at I2 | ND | ND | ND | ND | 2 |
| **0.3 msec** | All | 0.12 ± 0.1 | 8.22 ± 0.11 | 17.29 ± 0.17 | 37.07 ± 0.2 | 1207 |
| | Starting at folded | 0.07 ± 0.19 | 8.40 ± 0.21 | 17.45 ± 0.34 | 37.06 ± 0.44 | 288 |
| | Starting at I1 | --- | 8.10 ± 0.16 | 17.09 ± 0.32 | 37.10 ± 0.28 | 619 |
| | Starting at I2 | --- | --- | 17.38 ± 0.21 | 37.02 ± 0.41 | 299 |

8. Castillo JP, B. Tong A, Tafoya S, Jardine PJ, Bustamante C. A DNA packaging motor inchworms along one strand allowing it to adapt to alternative double-helical structures.

Nat Commun. 2021;12(1):1–8. doi:10.1038/s41467-021-23725-5 PubMed PMID: 34103515.

27. Humphrey W, Dalke A, Schulten K. VMD: Visual Molecular Dynamics. 1996.

28. Lindorff-Larsen K, Piana S, Palmo K, Maragakis P, Klepeis JL, Dror RO, et al. Improved side-chain torsion potentials for the Amber ff99SB protein force field. Proteins: Structure, Function and Bioinformatics. 2010 Jun;78(8):1950–8. doi:10.1002/prot.22711 PubMed PMID: 20408171.

29. Berendsen HJC, Grigera JR, Straatsma TP. The Missing Term in Effective Pair Potentials1. J. Phys. Chem [Internet]. 1987. Available from: https://pubs.acs.org/sharingguidelines
